# Pangenome Analysis of *Proteus mirabilis* Reveals Lineage-Specific Antimicrobial Resistance Profiles and Discordant Genotype-Phenotype Correlations

**DOI:** 10.1101/2025.11.21.689858

**Authors:** Namrata Deka, Aimee L. Brauer, Katherine Connerton, Blake M. Hanson, Jennifer N. Walker, Chelsie E. Armbruster

## Abstract

Urinary tract infections (UTIs) impose a large healthcare burden, with escalating antimicrobial resistance (AMR) and treatment failure. *Proteus mirabilis* is an under-characterized and challenging UTI pathogen due to intrinsic resistances and biofilm formation. To understand *P. mirabilis* population genomics, we combined pangenome analysis, *in silico* AMR predication, and phenotypic antimicrobial susceptibility testing (AST) across 1,027 *P. mirabilis* genomes derived from human urine specimens. This revealed a mosaic pangenome driven by extensive accessory genome plasticity. Multilocus sequence typing (MLST) identified 213 MLSTs with only 7% having ≥10 genomes, highlighting strain diversity. AMR gene profiles were largely lineage specific, with 25% of genomes harboring resistances for >6 antimicrobial subclasses. ST135 was identified as a highly MDR lineage, with 95% of genomes carrying ≥16 resistance genes. Mobile genetic element (MGE) analysis of 22 clinical isolates with complete, reference level genomes revealed that Tn7 transposons, IS26-mediated genomic islands, and class 1 integrons act as vehicles for high AMR gene dissemination, including IS26-mediated gene stacking within a *Proteus mirabilis* Genomic Resistance Island 1 (*PmGRI1)* in ST135 isolates. While presence of genes like *aph(3’)-la* reliably predicted kanamycin resistance, discordance for antibiotics such as trimethoprim-sulfamethoxazole and chloramphenicol revealed that AMR gene stacking, regulatory context, and intrinsic mechanisms, like efflux pumps, modulate phenotypic outcomes. In summary, our study provides a comprehensive and phenotypic resolution of *P. mirabilis* AMR, establishing that resistance architecture is lineage structured, MGE-driven, and phenotypically non-deterministic. We emphasize the need to shift towards standardized, genome-informed surveillance framework to translate into diagnostic and therapeutic strategies.

## INTRODUCTION

Urinary tract infections (UTIs) are among the most common bacterial infections, with an estimated healthcare cost of >$3 billion in the United States [1–4]. They are a leading cause of antimicrobial prescriptions in both outpatient and inpatient settings, with 50-60% of women experiencing at least one UTI in their lifetime [5]. UTI can be classified as complicated UTI (cUTI) or uncomplicated UTI (uUTIs) depending on the presence and absence of structural or functional abnormalities of the urinary tract, catheterization, or immunocompromised states [5]. Recent epidemiological data estimate approximately 174,000 cases of uUTI and 2.38 million cases of cUTI annually [6, 7]. Despite standard antibiotic therapy, treatment failure rates for UTIs remain prevalent, occurring in 16.7% of cases in adult female outpatients with cUTI, 27% of elderly men, and 15% of elderly women, often leading to recurrence, secondary infection, progression to pyelonephritis, or urosepsis [8, 9].

Among uropathogens, *Proteus mirabilis* is an important Gram-negative bacterium and a major cause of UTIs, contributing to approximately 4.2% of uUTIs and 4.7% of cUTIs [5]. Notably, *P. mirabilis* accounts for up to 45% of catheter-associated UTIs (CAUTIs) and 13-21% of secondary bacteremia, making it a key pathogen in medical device-associated infections [5, 10, 11]. *P. mirabilis* has a strong propensity for biofilm formation and causes encrustation of the catheter by deposition of struvite and apatite crystals, leading to persistent infections and catheter blockage [10, 12–15]. These urinary stones can remain in the bladder as crystalline deposits despite multiple doses of antibiotic treatment, catheter changes, and even periods without catheterization [15–17]. Infection with *P. mirabilis* can become fatal when it progresses to symptomatic CAUTI, acute bacteremia, and urosepsis due to its ability to disseminate from the bladder to other organs [14, 18].

Antimicrobial resistance (AMR) in UTI pathogens is an escalating global concern, with increasing reports of multidrug-resistant (MDR) strains complicating treatment [19]. Treatment failure for *P. mirabilis*-associated UTIs has been linked to MDR, leading to prolonged hospitalization, increased healthcare costs, and higher morbidity [15, 19, 20]. While antibiotic therapy remains the primary treatment approach, its efficacy is increasingly compromised by both intrinsic and acquired resistance mechanisms that limit therapeutic options [14, 21–23]. The resistance to first-line UTI antibiotics such as trimethoprim-sulfamethoxazole (TMP-SMX), fluoroquinolones, and β-lactams increases and the prevalence of ESBL genes like *blaTEM* and *blaCTX-M* further exacerbates resistance. Among the key drivers of AMR dissemination in *P. mirabilis* are mobile genetic elements (MGEs) such as SXT/R391 integrative and conjugative elements (ICEs) [19, 24, 25]. However, characterization of MGEs in *P. mirabilis* UTI clinical isolates remains limited. Understanding the MGE landscape is essential for deciphering the mechanisms underlying horizontal gene transfer, the evolution and spread of MDR, and the discordance between genotypic predictions and phenotypic resistance.

Numerous studies have characterized the model *P. mirabilis* strain HI4320, originally isolated from the female urinary tract, and its pathogenic potential [10, 26]. However, a major gap remains in our understanding of the diversity of strains isolated from recent clinical urine samples, either during infection or in the presence of a catheter. Multilocus sequence typing (MLST) provides an unambiguous method for characterizing bacterial isolates based on the sequences of internal fragments of typically 6-7 housekeeping genes. While previous studies in other bacterial species suggest that certain lineages correlate with varying levels of resistance and biofilm production, such associations in *P. mirabilis* remain largely unexplored. For example, in methicillin-resistant *Staphylococcus aureus* (MRSA), expression of the *ica* genes—critical for biofilm formation—varies among different STs, with certain STs showing higher *ica* gene expression, particularly in strains isolated from wound and catheter samples [27]. If similar correlations exist in *P. mirabilis*, routine sequence typing of clinical isolates could serve as a valuable tool to predict AMR potential and pathogenicity.

Several typing schemes have been developed to characterize *P. mirabilis* strains, but no standard scheme is followed for clinical classification. Core genome MLST (cgMLST), a high-resolution genomic method, has identified 205 clonal groups (CGs) linked to severe UTIs and AMR [28]. Historically, *P. mirabilis* strain differentiation has relied on methods such as bacteriophage typing, serotyping, and Dienes typing, but genomic approaches now offer superior discriminatory power [29–33]. Even with the established role of *P. mirabilis* in CAUTI, the publicly available genome database is limited compared to other UTI pathogens [32, 34, 35]. Furthermore, there remains a lack of integrated data linking sequence types (ST) to AMR gene carriage in urinary isolates using a standard classification framework. To address this gap, we focused specifically on genomes isolated from human urine, enabling a targeted analysis of AMR profiles and providing insights that may inform risk stratification for treatment failure and severe disease.

There are numerous ways to predict AMR, including the standardized clinical laboratory practices of testing resistance phenotypes to specific antibiotics, as well as genomic or machine learning methods to predict resistant gene expression [36]. Genomic prediction tools can be useful for quick detection of AMR in clinical practice, and there may be a correlation between AMR gene prediction and phenotypic expression, but the nature of the correlation can vary depending on specific genes and the bacterial species. For instance, *Escherichia coli* frequently harbors resistance genes for β-lactams, sulfonamides, trimethoprim, and chloramphenicol on mobile genetic elements, leading to a stronger correlation between genotype and phenotype [37–39]. In contrast, according to some studies, *Proteus mirabilis* carrying β-lactam and sulfonamide (*sul1* and *sul2*) genes exhibits this correlation less consistently, showing weaker alignment between genomic predictions and observed resistance profiles [37, 40–42].

Tools like AMRFinder and CARD are accurate *in silico* prediction tools; AMRFinder was able to validate the presence of genes pertaining to AMR phenotypes for *Salmonella enterica, Campylobacter spp*., and *E. coli* with 98.4% consistent predictions [43]. Further, a 99% concordance was found between phenotypic antimicrobial susceptibility testing and *in silico* prediction of AMR genes using whole-genome sequencing (WGS) in 1,321 isolates of *Salmonella enterica* from human infections in Canada [20]. While WGS-based methods provide a promising avenue to predict AMR in other species, no equivalent variation exist for *P. mirabilis*. Hence, integrating genotypic AMR prediction and phenotypic MIC testing in diverse strains may offer a more comprehensive approach to predicting antimicrobial resistance in this species.

In this study, we conducted a comparative genomic analysis of *P. mirabilis* using 1,001 publicly available human urine isolate sequences to characterize the pangenome, AMR gene profiling, and *in-silico* MLST typing. We benchmarked this against 26 clinical isolates collected from patients with long-term indwelling catheters [15, 17]. Our analysis revealed that AMR gene profiles can link to specific lineages rather than isolation source, with isolates of the same MLST clustering together and exhibiting related resistant genes. We found that ST135 lineage is a hotspot for resistance gene acquisition in our dataset, driven by the integration of MGEs such as IS26 and the *PmGRI1* genomic island with multiple overlapping resistance genes. We further identified a high rate of discordance between genotype-based susceptibility prediction and phenotypic resistance. This discrepancy can be partially attributed to uncharacterized MGE’s and efflux pump mechanisms, highlighting the limitations of relying on gene detection only for resistance profiling. Our findings underscore the need for integrating lineage-specific MGE surveillance with phenotyping testing to guide treatment options and reduce the overuse of broad-spectrum antibiotics in managing *P. mirabilis* CAUTI infections.

## MATERIALS AND METHODS

### Data availability statement

Sequencing data have been deposited in NCBI BioProject PRJNA1367153: *Proteus mirabilis* genome sequences from human urine sources in the United States. All other data are available within the article and its supplemental material. All basic commands and databases used are available in the supplementary appendix.

### Ethics approval

This body of work includes sequencing of *P. mirabilis* strains that were previously isolated from human subjects in two different studies, for which human subject protocols were approved by the University of Buffalo (IRB #STUDY00002526) and Washington University School of Medicine (IRB #201410058).

### Patient population and Sample collection

To investigate whether *P. mirabilis* MLSTs correlate with AMR patterns, we assessed *P. mirabilis* genomes from three separate cohorts: publicly available genomes from NCBI (**Table S1**), nursing home residents with long-term catheters (CEA isolates), and community-dwelling catheterized patients (HUC isolates).

For comparison of *P. mirabilis* genomes, we sought to query all publicly available high-quality genomes of *P. mirabilis* isolates obtained from urine samples in hospital environments in the United States. Using advanced search parameters, publicly available raw genome data were retrieved from the NCBI database. The initial search, conducted in October 2024, yielded 1,337 hits using the terms “*Proteus mirabilis*,” isolation source “urine,” and host “*Homo sapiens*.” Filtering the results to include only isolates originating from the “United States” reduced the dataset to 1,152 genomes. The dataset was further refined by only including whole-genome sequencing data. These filters resulted in a final dataset of 1,064 (**Table S1**) genomes, which were subsequently downloaded from NCBI. Following FastQC v0.11.9-Java-11 analysis and adapter trimming using trimmomatic v0.39-Java-11.0.16. [44, 45], 1,026 genomes were retained for downstream analyses. Samples were excluded if they exhibited poor quality, had high GC content, high read duplication rates, or if paired end read files were missing (46 samples). Raw paired end read sequences were assembled using spades v3.15.5 [46]. After genome assembly, we assessed quality and completeness using QUAST (**Table S2**), selecting 1,001 high-quality genomes with >95% completeness and fewer than 1,000 contigs which were annotated with prokka v1.14.5 to facilitate downstream analyses [47, 48].

Isolates designated 101–210 were cultured from urine specimens collected during a prospective cohort study on asymptomatic catheter-associated bacteriuria at two nursing homes in Buffalo, New York, USA, between July 2019 and March 2020 [15]. Inclusion criteria required participants to be 21 years or older at the time of consent and have an indwelling urinary catheter in place for at least 12 months at the start of the study. Study participants underwent an initial enrollment visit, followed by weekly study visits for up to seven months. Most of the study participants were White (79%) and male (79%), an average age of 65 years, and 68% had suprapubic catheters [15]. Participants exhibited a high degree of functional dependence in activities of daily living, with the most common comorbidities including neurogenic bladder (74%), hemiplegia (42%), diabetes (32%), renal disease (32%), multiple sclerosis (26%), and chronic heart failure (26%) [15]. Urine specimens were cultured on MacConkey agar and Columbia Nutrient Agar, and Gram-negative bacteria were identified to the species level using API-20E test strips (BioMérieux). *P. mirabilis* species designation was further confirmed based on characteristic swarming motility on blood agar plates (Hardy Diagnostics) and whole genome sequencing (**Table S3**).

Additional *P. mirabilis* isolates with designations of “HUC” were obtained from the lab of Scott J. Hultgren, Washington University School of Medicine [17]. These isolates were derived from urine samples collected from community-dwelling catheterized patients during routine catheter removals in the urology clinic as part of their standard care. Most of the participants were male (64%) with an average age of 64 years. Inclusion criteria required participants to be 18 years or older at the time of consent and scheduled for the removal of an indwelling urinary device, including urethral and suprapubic catheters or ureteral stents. The primary indications for catheterization included bladder outlet obstruction (33%), peripheral or central neuropathy (40%), rectovesical or rectourethral fistula (7%), stress urinary incontinence (7%), atonic bladder (11%), and temporary post-operative complications (2%) [17]. Bacterial colonies were distinguished based on morphology, size, and color, and 1–9 single colonies per sample were selected for DNA extraction. *P. mirabilis* isolates were identified via 16S rRNA sequencing and confirmed by whole genome sequencing (**Table S3**).

### Whole Genome Sequencing of Clinical Isolates

Freezer stocks were streak-plated on MacConkey agar to isolate single colonies of *Proteus mirabilis.* 5 ml of Low Salt Luria Bertani (LSLB; 0.1 g/L NaCl, 10 g/L Tryptone, 5 g/ yeast extract) broth was inoculated with a single colony and incubated for 16 hours at 37°C with shaking. Genomic DNA was extracted using the DNeasy Blood and Tissue Kit (Qiagen, Germantown, Maryland, USA) according to the manufacturer’s instructions with specific modifications to ensure high purity of DNA; samples were incubated in proteinase K for 2 hours instead of 10 minutes, and three washes of AW2 buffer were conducted to ensure high-quality DNA free of salts. Samples were sent to SeqCoast Genomics for whole genome sequencing (WGS). All samples were prepared using the Illumina DNA Prep tagmentation kit and IDT For Illumina Unique Dual Indexes.

Sequencing was performed on the Illumina NextSeq2000 platform using a 300-cycle flow cell kit to produce 2×150bp paired reads, as previously described [49]. 1-2% PhiX control was spiked into the run to support optimal base calling. The sequencing data has been uploaded in BioProject PRJNA1367153: *Proteus mirabilis* genome sequences from human urine sources in the United States. Read trimming and run analytics were performed using trimmomatic v0.39.0 [45]. For long read sequencing, Oxford Nanopore Technologies Native Barcoding Kit (#SQK-NBD114) and long fragment buffer was used to promote longer read lengths of DNA. DNA was sequenced on the PromethION 2 Solo platform using a FLO-PRO114M Flow Cell (vR10.4.1). Sequencing was performed with a translocation speed of 400bps. Base calling was done on the GridION using the super-accurate basecalling model with barcode trimming enabled.

Assembly of short reads from the clinical isolates and the genomes acquired from the NCBI database was performed using spades v3.15.5, while hybrid assembly of long and short reads from clinical isolates was performed using Unicycler v0.5.0 [46]. Here, long reads were used to assemble the genome structure and the short reads were used to polish the genome. The assemblies were quality-checked using QUAST v5.2.0 selecting genomes with < 500 contigs for only short read assembled genomes and <3 contigs for hybrid (**Table S3**) If sequences were not of good quality, they were further polished using pilon/1.23-Java-11.0.16 [50]. Genomes that passed quality filtering were annotated using prokka v1.14.5 [48].

### Multi Locus Sequence Typing (MLST) and Phylogenetic trees

Multilocus sequence typing (MLST) analysis was conducted to identify the sequence types (STs) present in the dataset using mlst software, which incorporates the PubMLST database [51, 52]. The annotated genomes were then fed into roary v3.13.0 using a core threshold cluster of 99%, to identify their pangenome profile and develop a gene presence/absence matrix [53]. The filtered core genome alignment was constructed using FastTree v2.1.11. The resulting newick file was visualized as a mid-rooted phylogenetic tree using the iTOL website and annotated with source, and antibiotic metadata files [54, 55].

### Drug Resistance Gene Prediction

AMRFinderPlus (using the Bacterial Antimicrobial Resistance Reference Gene Database) and CARD (The Comprehensive Antibiotic Resistance Database) were used to predict AMR genes from the coding sequence of the *P. mirabilis* genomes including the 26 CEA and HUC strains and clinical reference strain HI4320 [36, 43].

Mobile genetic elements were characterized in the hybrid-sequenced clinical isolates using multiple approaches. Insertion sequence (IS) elements were identified using ISEScan v1.7.3 [56]. Prophage regions were predicted using PHASTER (PHAge Search Tool Enhanced Release) [57–59] and classified as intact (score >90), questionable (70-90), or incomplete (<70). Plasmid replicons were detected using ABRicate v1.0.1 with the PlasmidFinder database (≥80% identity, ≥60% coverage) [60, 61]. Integrative and conjugative elements (ICEs) were identified via ICEberg 3.0 [62]. Spearman’s rank correlation in R v4.3.0 assessed associations between IS abundance and AMR gene content and virulence factors with significance set at p <0.05.

### Antibiotic Susceptibility Testing (AST)

The clinical strains were grown overnight in 5 mL LSLB broth at 37°C with shaking. Antibiotic solutions: Tetracycline HCl (Research Products International (RPI), CAS: 64-75-5), Chloramphenicol (RPI, CAS: 56-75-7), Streptomycin (RPI, CAS: 3810-74-0), Streptothricin (Goldbio, CAS: 96736-11-7), Sulfonamide (sulfamethoxazole, Cayman chemical company, CAS: 723-46-6), Trimethoprim (Cayman chemical company, CAS: 738-70-5), Ampicillin (RPI, CAS: 69-53-4), and Kanamycin (RPI, CAS: 8063-07-8), were prepared in Mueller-Hinton Broth (MHB) at a concentration range encompassing the sensitive-to-resistant spectrum according to Clinical Laboratory Standards Institute (CLSI) guidelines for *Enterobacteriales* [63]. Stock solutions of all the antibiotics were prepared and stored at −20°C, and working dilutions were made fresh from the stock immediately prior to testing.

1 ml of the overnight cultures were centrifuged at 1,000g to pellet the cells. The supernatant was removed, and the pellets were resuspended in 0.9% saline to wash away residual LB. The inoculum was adjusted to 4×10^5^ CFU/ml in MHB. One antibiotic with increasing concentrations was tested per 96-well plate, with four technical and duplicate biological replicates. The plates were incubated at 37°C with double-orbital shaking in Biotech Synergy H1, and the OD_600_ was measured every 15 minutes for 20 hours to monitor bacterial growth with OD_600_ threshold set at 0.15.

### Statistical analysis

Logistic regression models were used to explore AMR gene carriage as a function of sequence type, geographic location of isolation, and year of isolation. Data were analyzed using Stata/IC version 15.1 (StataCorp LP, College Station, TX).

## RESULTS

To assess the population structure and genetic diversity of *Proteus mirabilis* in the urinary tract, we analyzed 1,001 publicly available genomes isolated from human urine specimens (**Table S1**). The genomes downloaded from NCBI were predominantly derived from clinical settings, focusing on medical surveillance and characterization of urinary samples.

Key contributing sources include research from Washington University School of Medicine (WUSM), the University of Pittsburgh Enhanced Detection System for Healthcare-Associated Transmission (EDS-HAT) study, and national antimicrobial surveillance programs such as the Center of Disease Control’s Healthcare-Associated Infection Sequencing (CDC HAI-Seq) Gram-Negative Bacteria Study and studies on the human urinary microbiome. Among the genomes, contig counts ranged from 60 to 973, with an average of 258 and a median of 180 contigs. The GC content averaged 38.76%, with a median of 38.69%, making them suitable for downstream analyses (**Table 1 and S2**). While all sequences were confirmed to be derived from urine samples, the clinical context of these specimens—whether they were associated with asymptomatic bacteriuria or infection—is not known.

**Table 1:** Summary statistics of *P. mirabilis* genome sequences.

| Publicly available *P. mirabilis* genomes isolated from urine |  |  |  |  |
| --- | --- | --- | --- | --- |
| Metric | Average | Max | Min | Median |
| # Contigs | 258.43 | 973 | 60 | 180 |
| Total Length | 4020761.51 | 8382181 | 3667799 | 3992140 |
| N50 | 174294.28 | 1994055 | 4981 | 174571 |
| GC Content | 38.76 | 43.24 | 37.57 | 38.69 |
| Clinical *P. mirabilis* genomes isolated from catheterized individuals |  |  |  |  |
| Metric | Average | Max | Min | Median |
| # Contigs | 66.52 | 98 | 2 | 69 |
| Total Length | 4015048.48 | 4165371 | 3795264 | 4015495 |
| N50 | 317358.93 | 4063606 | 119871 | 182416 |
| GC Content | 38.74 | 38.98 | 38.38 | 38.78 |

Phylogenetic analyses (mid-rooted tree) revealed a monophyletic, star-like phylogeny, with a central ancestral node from which several peripheral branches radiate (**Fig 1a**), which may indicate recent diversification events. When considering branching, a subset of genomes branched out creating three clades, *clade 1* (964 genomes, dominant), *clade 2* (5 genomes), and *clade 3* (32 genomes) (**Fig S1**). This breakdown is consistent with previously reported intra-species core genome variation in *P. mirabilis* [32, 33, 35]. For example, Potter *et. al.* studied a cohort of 2,060 *P. mirabilis* genomes collected worldwide from multiple anatomical sources and observed that the essential core genes remained relatively conserved with three divergent sub-species. [32].

**Fig 1.**
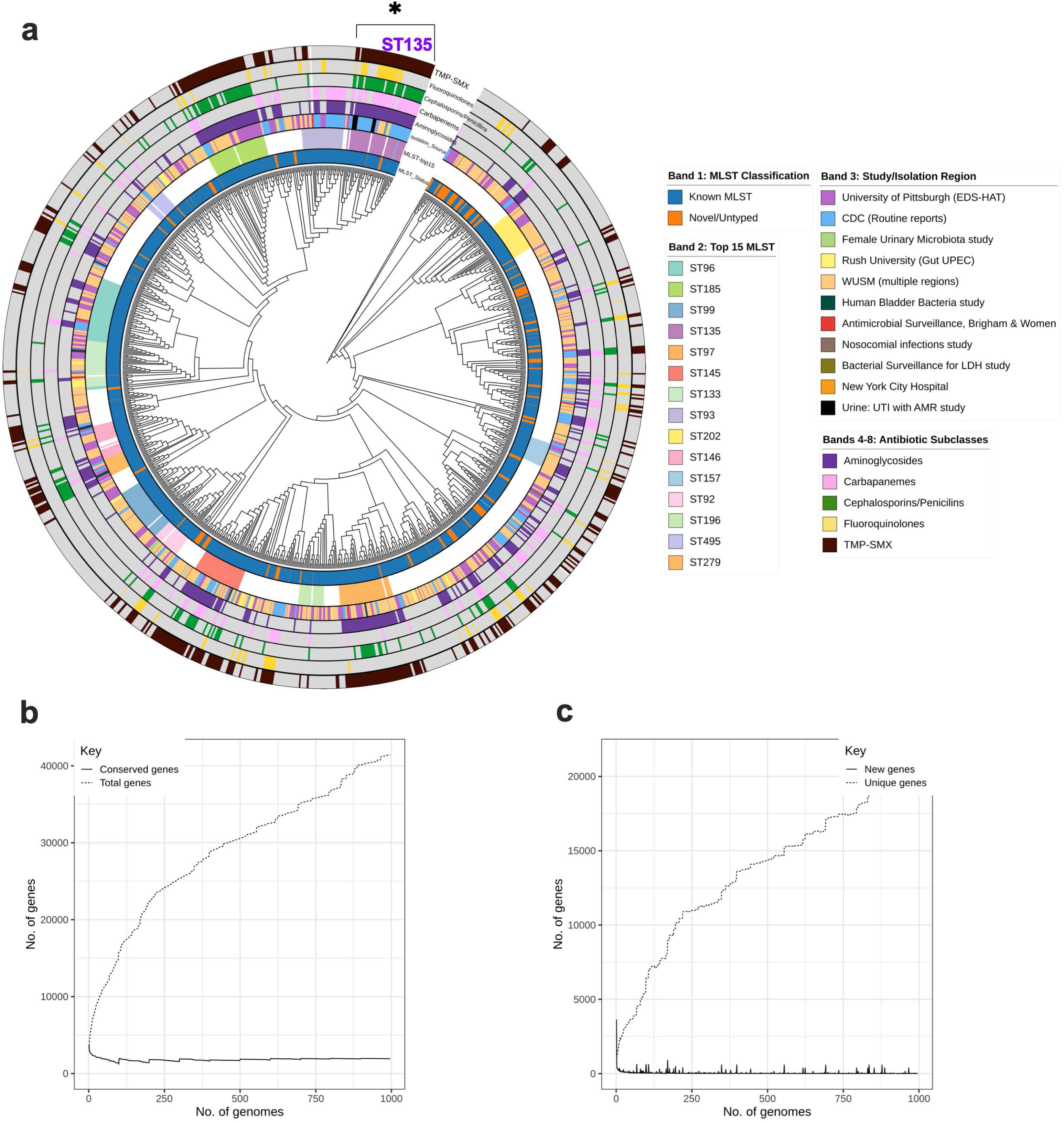
Phylogenetic tree and pangenome analysis of publicly-available *Proteus mirabilis* genomes from human urine samples. (**1a**) maximum-likelihood phylogenetic tree of 1,001 *P. mirabilis* human urine genomes with metadata arranged as concentric bands. The first two bands display multi-locus sequence type (ST), and the third band indicates the study source in the NCBI database associated with the Bioproject. Bands 4-8 indicate presence (colored bars) or absence (grey) of resistance genes across different antibiotic classes. Distinct clusters were observed that either had resistance genes for nearly all classes of antibiotics or lacked them entirely, which was dependent on ST and independent of geographical source. (**1b**) Roary collector curve showing conserved genes (∼1,700 core genome) present in all of the genomes analyzed as well as the number of total genes, which increases as new genomes are added, indicating an open pangenome. (**1c**) Roary collector curve showing unique vs new gene accumulation across all *P. mirabilis* genomes. Here, the total unique genes is >15,000 indicating massive accessory gene diversity and an open pangenome. Early genomes introduce a lot of new genes before dropping out. However, after addition of >250 genomes, new genes (vertical lines) are still accumulated, indicating horizontal gene transfer (HGT), mobile genetic elements, or prophage insertions.

### Pan-genomic analysis of *P. mirabilis* reveals a mosaic structure

To characterize the pangenome of *P. mirabilis* from human urine samples we assessed its structural composition, including core, shell, and cloud gene categories. In the 1,001 genomes analyzed, a total of 41,121 gene families were identified. Of these, the core genome (genes conserved in >99% of the samples analyzed) is small with 1,937 genes indicating a high degree of conservation. Considering that a typical *P. mirabilis* genome harbors roughly 4,600-4,900 coding sequences (CDS), the core genome accounts for approximately 40-45% of the average gene content. This indicates a highly conserved genomic backbone with 4.7% of it supporting essential cellular functions, coexisting with an expansive and dynamic accessory genome (**Fig S1**). The shell of the 1,001 genomes contained 1,749 genes (4.08%), and 531 soft core genes shared among a subset of strains that likely contribute to niche-specific adaptations. The majority of genes (37,204 genes, 90%) were classified as cloud genes, making the accessory component the largest in the pangenome. Having a small, stable core and large, variable accessory genome likely represents strain-specific differences driven by horizontal gene transfer.

The pangenome exhibited an open conformation representing a mosaic pattern (**Fig S1**), with the size expanding as additional genomes were analyzed (**Fig 1b**). This expansion reflects the continuous acquisition of novel gene families, highlighting the dynamic nature of the *P. mirabilis* genome and its capacity for adaptation through horizontal gene transfer (**Fig 1c**). The stability of core genes is crucial for the consistent expression of virulence factors, while the expanding accessory genes may help adapt to polymicrobial environments and/or antibiotic pressures. These findings indicate the genetic plasticity of *P. mirabilis* isolated from urine and provide a foundation for future studies on its pathogenic potential and AMR mechanisms in the urinary tract.

### Publicly available *P. mirabilis* genomes comprise 213 defined and 118 undefined MLSTs

The specific genes used in MLST of *P. mirabilis* are a combination of housekeeping genes *atpA, dnaJ, mdh, pyrC, recA,* and *rpoD* [51, 52]. We identified a total of 213 defined STs and approximately 118 novel or undefined genomes, reflecting the diversity of *P*. *mirabilis* human urine isolates within the dataset (**Fig 1**). The untyped MLST have been submitted to the PubMLST database to update the schema in *P. mirabilis*.

A detailed analysis of ST prevalence within the dataset revealed that certain STs were more frequently observed. The distribution of MLSTs across the dataset was highly skewed (**Fig 2a, Table S4 and S5**), with 46.26% of the STs having only one representative genome in the dataset while ∼7% of the STs had ≥10 genomes (**Fig 2a**). Many genomes in the same ST came from the same study/location but it was not the rule as the top 15 overrepresented ST’s show mixed isolation sources (**Fig 1a**). The uneven ST distribution is further visualized in a bar chart of the top 20 MLSTs, highlighting the dominance of a few clonal lineages across sampled populations (**Fig 2b**). ST96 was the most abundant (49 genomes; 4.90%), followed by ST185 (42 genomes; 4.20%), ST99 (38 genomes; 3.80%), ST135 (38 genomes; 3.80%), and ST97 (37 genomes; 3.70%). As expected, genomes from each of these STs clustered together by lineages, even if isolated from different geographical locations and reported in different studies (**Fig 1a**).

**Fig 2.**
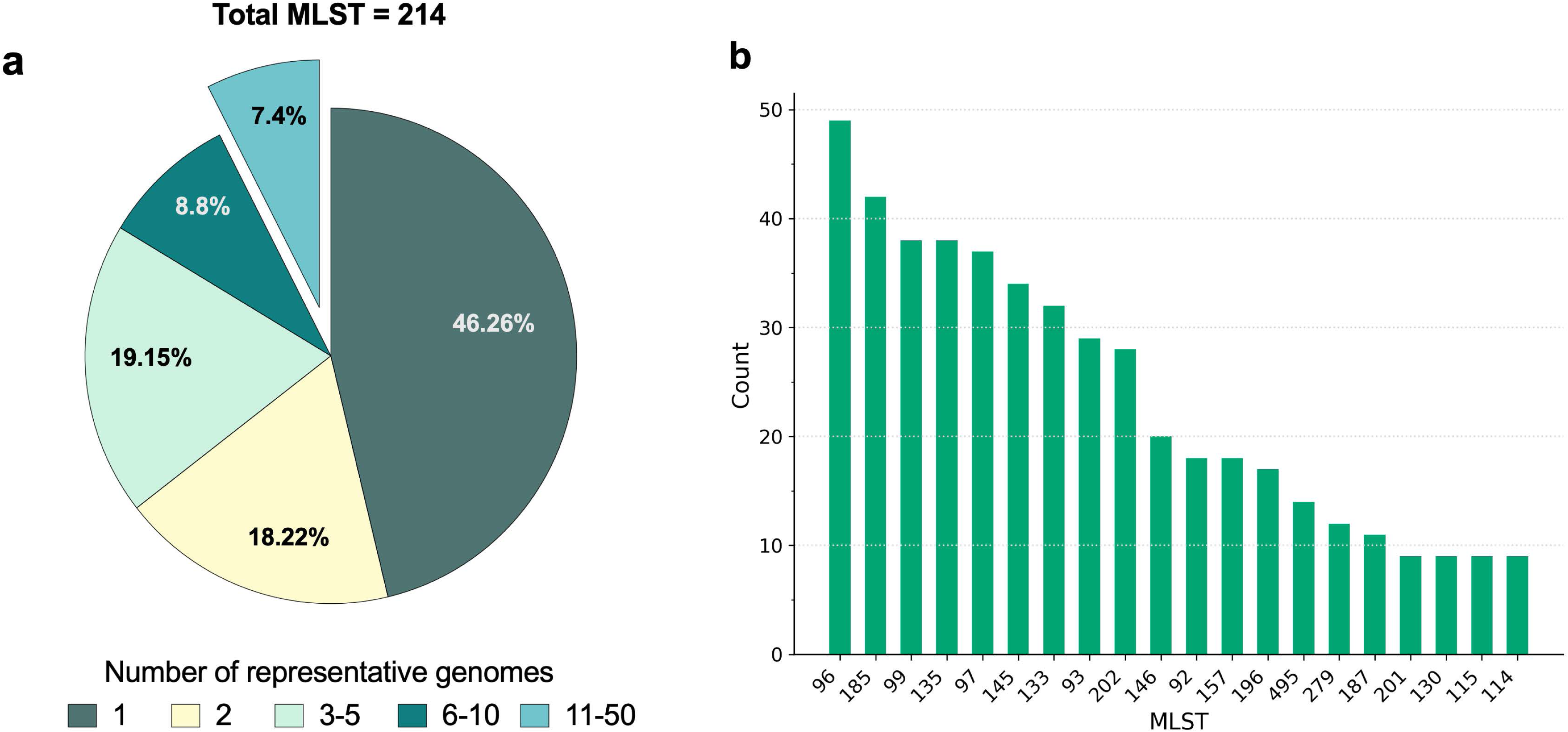
Sequence type distribution of publicly-available *Proteus mirabilis* genomes from human urine samples. (**2a**) Pie-chart illustrating the distribution of MLSTs grouped by their sample count frequency. (**2b**) Bar plot showing the top 20 MLSTs based on sample count.

### Phylogenetic analysis reveals shared lineages and novel variants across clinical ***P. mirabilis* isolates**

To further examine the clinical relevance of *P. mirabilis* MLSTs, we next sequenced and analyzed a collection of 25 clinical isolates from two distinct cohorts (CEA isolates collected from the urine of nursing home residents with long-term indwelling catheters, and HUC isolates collected from the urine of catheterized community-dwelling individuals during routine catheter changes), as well as the reference strain HI4320 that was previously isolated from the urine of a catheterized nursing home resident (**Fig 3a**). The analysis revealed a population structure forming two primary clades, termed *clade 1* (23 isolates) and *clade 2* (3 isolates) (**Fig 3b**). Lack of the third clade identified in the 1,001 genomes was expected considering its low frequency (0.5%) and the small sample size of the clinical isolate collection.

**Fig 3:**
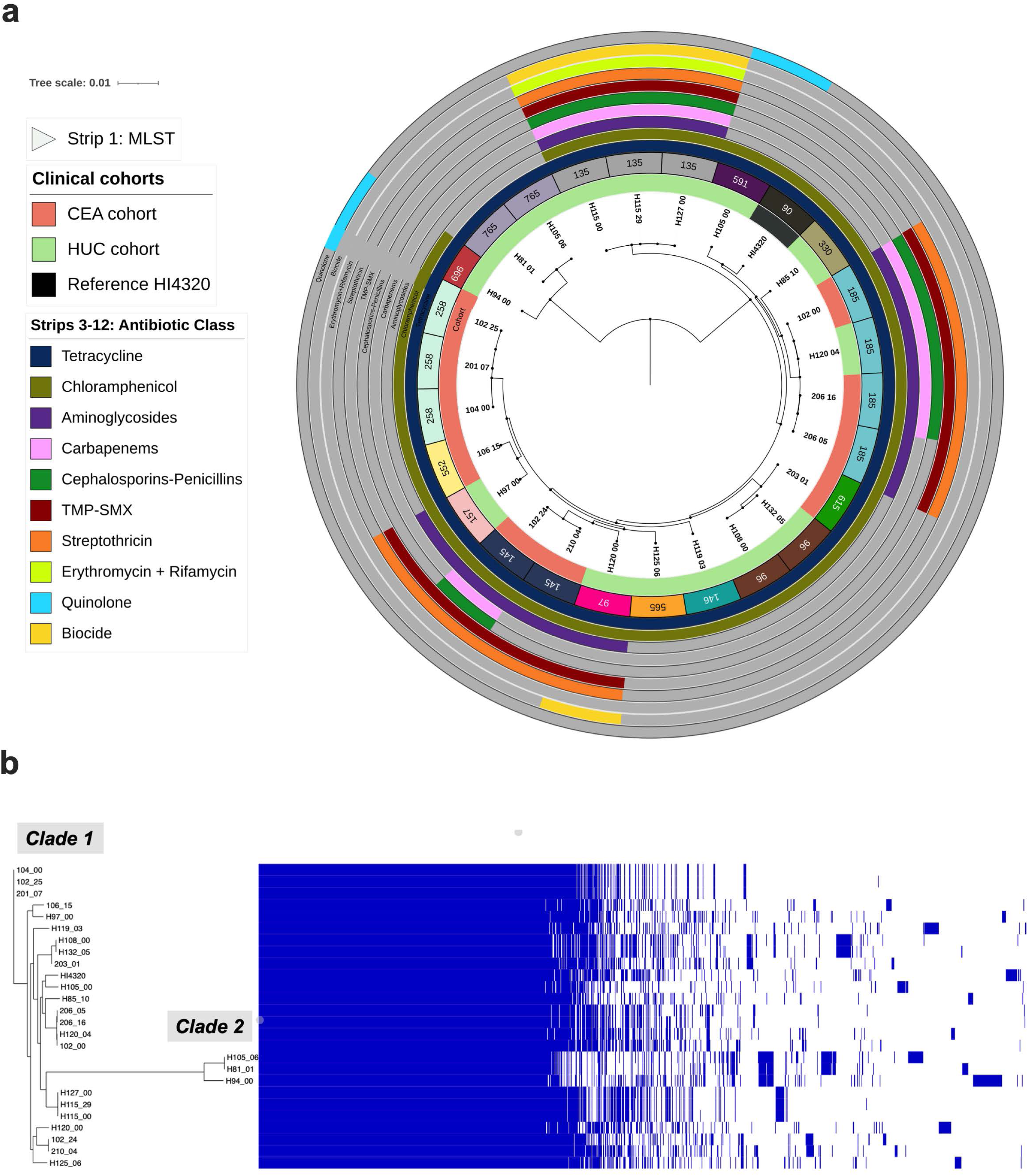
Phylogenetic tree and pangenome analysis of *Proteus mirabilis* strains isolated from individuals with indwelling urinary catheters. (**3a**) A mid-rooted maximum likelihood phylogenetic tree of clinical *P. mirabilis* isolates collected from patients with long-term urinary catheters. The phylogenetic tree displays the participant ID, the cohort ID (strip1), and multi-locus sequence type (ST, strip 2). Many of the isolates cluster by ST, but not by geographical location. (**3b**). Mosaic pangenome with presence/absence matrix of core-genome phylogenetic tree for 26 *P. mirabilis* clinical isolates collected from patients with long-term urinary catheters.

Phylogenetic clustering was in accordance with the core genome and gene content irrespective of the clinical origin (HUC vs CEA, **Fig 3a**). Across the 26 isolates, we identified 15 distinct STs; 10 STs were identified in the HUC cohort, 4 in the CEA cohort, and 1 represented solely by HI4320. Six of the 10 overrepresented STs in the NCBI genome dataset were also represented among the clinical isolates; namely STs 96, 97, 135, 145, 146, and 185. ST185 was observed in both the CEA and HUC cohorts (H120-04 and 102-00, 206-16), demonstrating genetic overlap across different geographic locations. The two most prevalent STs in the NCBI dataset were also the most prevalent in the clinical isolates: ST185 (3 isolates, 12%) and ST96 (3 isolates, 12%). Their presence confirms that our clinical cohort captured high-impact, epidemiologically relevant STs rather than rare or sporadically enriched variants even with a limited sample size. The full summary of isolated sources and STs are provided in **Table 2 and Table S3.**

**Table 2:** *P. mirabilis* clinical isolates used in this study.

| **Isolate ID** | **Study** | **Species** | **MLST** | **Patient designation in cited publication** | **Isolation week** |
| --- | --- | --- | --- | --- | --- |
| 102-00 | CEA | *P. mirabilis* | 185 | Participant A [15] | Visit 0 |
| 102-24 | CEA | *P. mirabilis* | 145 | Participant A [15] | Visit 24 |
| 102-25 | CEA | *P. mirabilis* | 258 | Participant A [15] | Visit 25 |
| 104-00 | CEA | *P. mirabilis* | 258 | Participant C [15] | Visit 0 |
| 106-15 | CEA | *P. mirabilis* | 552 | Participant H [15] | Visit 15 |
| 201-07 | CEA | *P. mirabilis* | 258 | Participant G [15] | Visit 7 |
| 203-01 | CEA | *P. mirabilis* | 615 | Participant F [15] | Visit 1 |
| 206-05 | CEA | *P. mirabilis* | 185 | Participant D [15] | Visit 5 |
| 206-16 | CEA | *P. mirabilis* | 185 | Participant D [15] | Visit 16 |
| 210-04 | CEA | *P. mirabilis* | 145 | Participant J [15] | Visit 4 |
| H81-01 | HUC | *P. mirabilis* | 765 | Patient 81-urine [17] | Visit 1 |
| H85-10 | HUC | *P. mirabilis* | 330 | Patient 85-urine [17] | Visit 10 |
| H94-00 | HUC | *P. mirabilis* | 696 | Patient 94-urine [17] | Visit 0 |
| H97-00 | HUC | *P. mirabilis* | 157 | Patient 97-urine [17] | Visit 0 |
| H97-11 | HUC | *P. mirabilis* | 157 | Patient 97-urine [17] | Visit 11 |
| H105-00 | HUC | *P. mirabilis* | 591 | Patient 105-urine [17] | Visit 0 |
| H105-06 | HUC | *P. mirabilis* | 765 | Patient 105-urine [17] | Visit 6 |
| H108-00 | HUC | *P. mirabilis* | 96 | Patient 108-urine [17] | Visit 0 |
| H115-00 | HUC | *P. mirabilis* | 135 | Patient 115-urine [17] | Visit 0 |
| H115-05 | HUC | *P. mirabilis* | 135 | Patient 115-urine [17] | Visit 5 |
| H119-03 | HUC | *P. mirabilis* | 146 | Patient 119-urine [17] | Visit 3 |
| H120-00 | HUC | *P. mirabilis* | 97 | Patient 120-urine [17] | Visit 0 |
| H120-04 | HUC | *P. mirabilis* | 185 | Patient 120-urine [17] | Visit 4 |
| H125-06 | HUC | *P. mirabilis* | 565 | Patient 125-urine [17] | Visit 6 |
| H127-00 | HUC | *P. mirabilis* | 96 | Patient 127-urine [17] | Visit 0 |
| H132-05 | HUC | *P. mirabilis* | 96 | Patient 132-urine [17] | Visit 5 |
| HI4320 | Reference | *P. mirabilis* | 90 | Refseq:<br>GCA_000069965.1_A<br>SM6996v1_genomic | - |

### *Proteus mirabilis* urine isolates possess resistance factors to multiple subclasses of antibiotics

To understand how AMR complicates *P. mirabilis* UTI treatment and contributes to the global public health burden, we examined the AMR profiles of the 1,001 publicly available *P. mirabilis* genomes. A total of 24 antibiotic resistance subclasses were identified, reflecting the diverse mechanisms by which this pathogen can evade antibiotics. The most prevalent genes were those conferring resistance to tetracycline and chloramphenicol, observed in nearly all genomes (**Table S6)**.

*P. mirabilis* is known to have intrinsic resistance to tetracycline, so it was not surprising that tetracycline resistance genes were detected in 991 (99.0%) of the 1,001 genomes, with *tetJ* specifically encoded by 985 of those 991 (99.5%) (**Table 3**). Several other tetracycline resistance determinants, including *tetA, tetB, tetC*, and *tetD*, were detected at lower frequencies. Interestingly, 934/991 (94.2%) of the genomes carried only a single *tet* gene, while 48 (4.8%) carried 2, and 9 (0.9%) carried 3 *tet* genes (**Fig 4a**). Notably, a small number of genomes (11, 1.1%) had no detectable tetracycline resistance genes. This could represent a true absence of the genes, or it could be a result of fragmented assemblies or sequence dissimilarity not found in the AMR prediction database.

**Fig 4:**
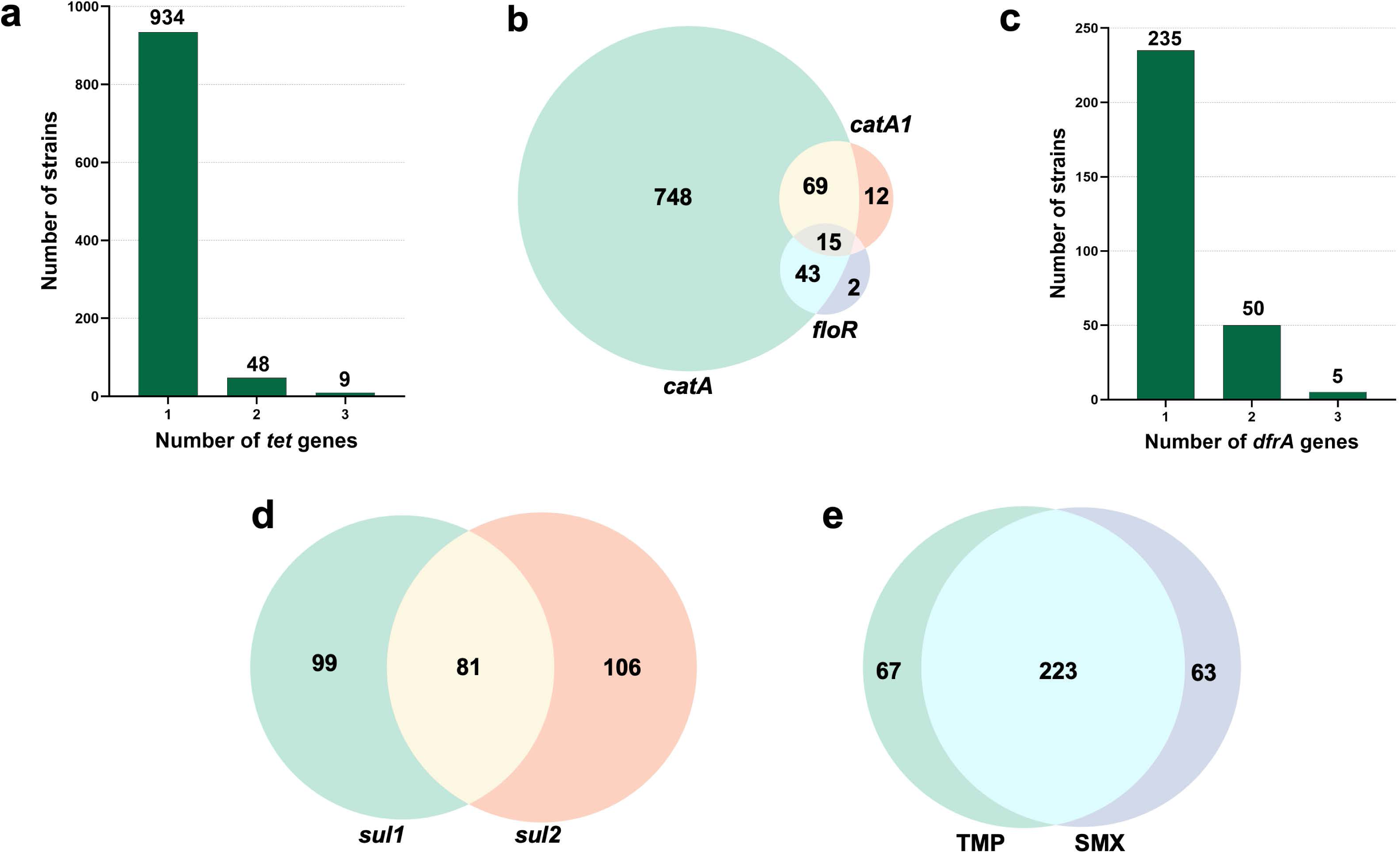
AMR gene carriage in publicly-available *P. mirabilis* genomes. (**4a**) Number of *P. mirabilis* strains in the NCBI dataset carrying 1, 2, and 3 tetracycline resistance (*tet)* genes. (**4b**) Three-way Venn diagram exhibiting the shared chloramphenicol resistance genes (*catA, catA1,* and *floR*) by all the *P. mirabilis* isolates in the NCBI dataset. (**4c**) Number of *P. mirabilis* strains in the NCBI dataset carrying 1, 2, and 3 trimethoprim resistance (*dfrA)* genes. (**4d**) Venn diagram of shared sulfonamide resistance genes (*sul1* and *sul2*). (**4e**) Venn diagram of shared trimethoprim (TMP) and sulfonamide (SMX) resistance genes.

**Table 3:**
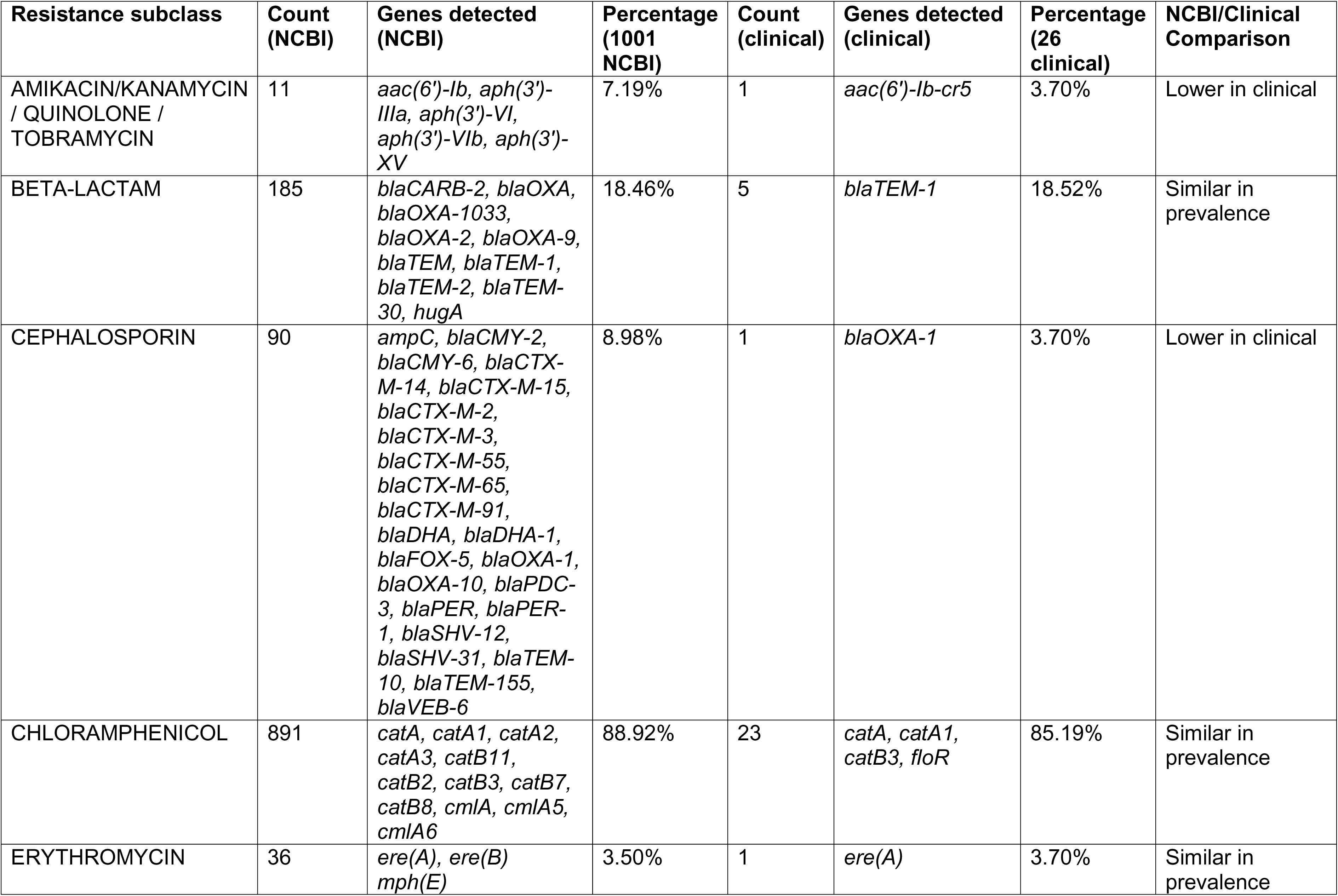
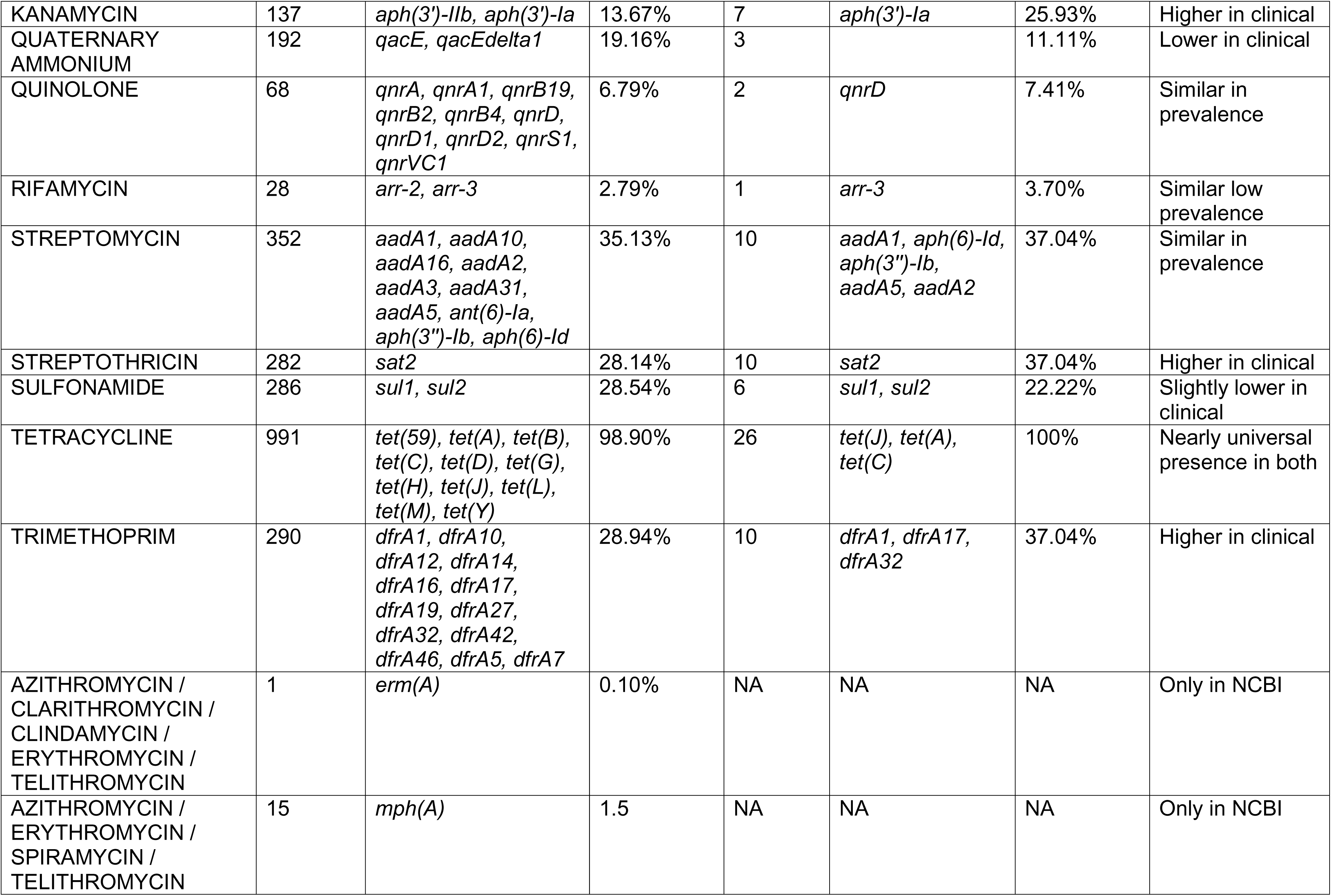
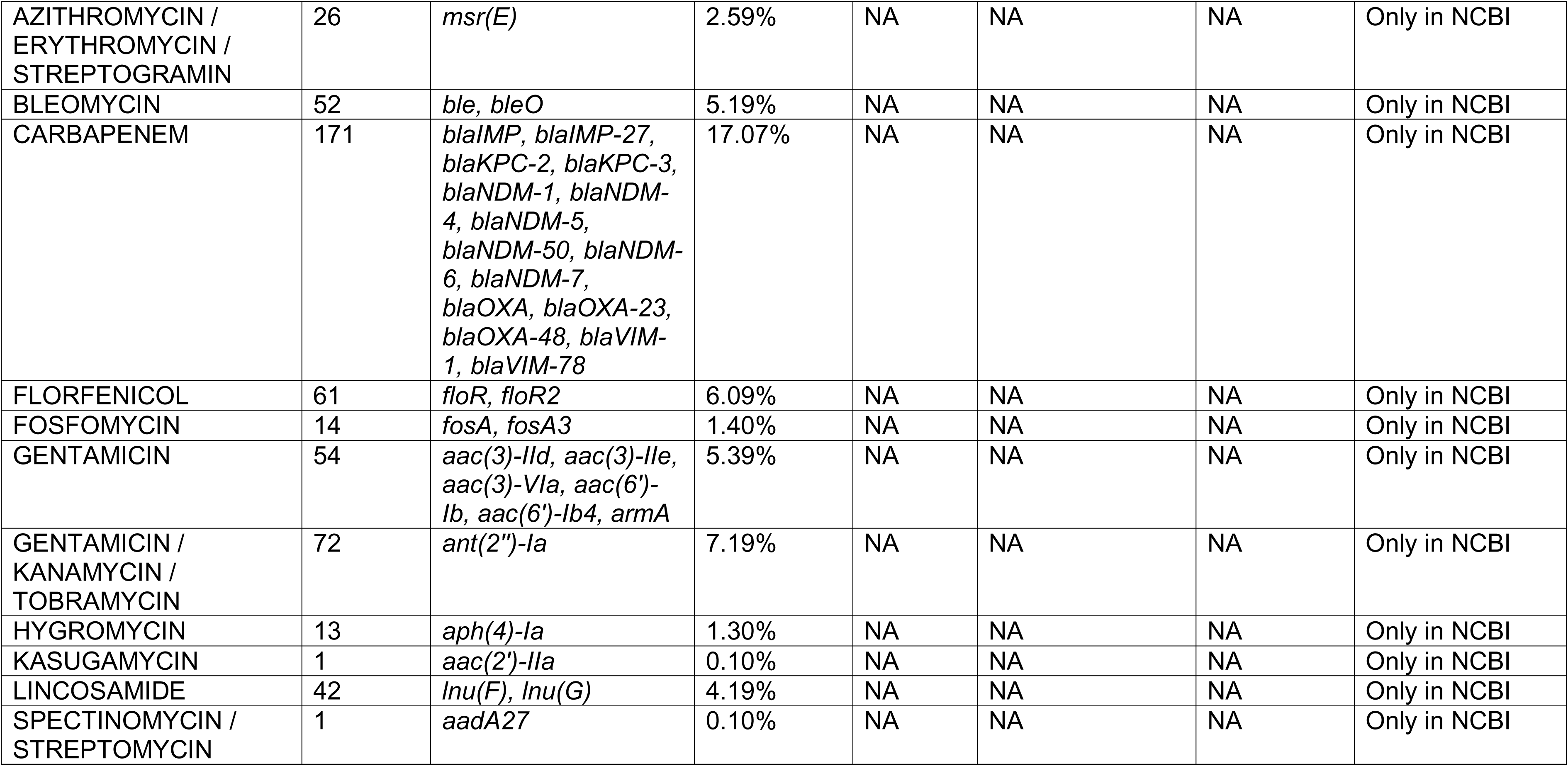
AMR subclass count and percentage in *P. mirabilis* genomes.

Chloramphenicol resistance was detected in 952 (95%) of the genomes, with 748 harboring the *catA* gene. Additionally, 69 genomes harbored both *catA* and *catA1,* 43 harbored *catA* and *floR,* and the presence of all three genes was seen in 15 genomes (**Fig 4b, Table S6**). These findings highlight the prevalence of phenicol resistance in *P. mirabilis* genomes despite limited use of chloramphenicol in modern clinical settings.

Resistance to aminoglycosides, a critical class of antibiotics for treating severe Gram-negative infections, was predicted in 716 (71%) of the genomes. Notably, we observed streptomycin resistance genes in 352 genomes, suggesting the persistence of resistance despite limited clinical use (**Table 3**). The presence of multiple aminoglycoside resistance determinants indicates that *P. mirabilis* maintains a robust arsenal against this class of antimicrobials, including amikacin, gentamycin, kanamycin, streptomycin and tobramycin.

Resistance to trimethoprim (TMP), mediated by *dfrA* alleles, was detected in 290 (29%) genomes. The most prevalent allele was *dfrA1* (n=248), followed by *dfrA17* (n=44), and *dfrA32* (n=17). 235 (81.02%) genomes had a single *dfrA* gene, while 50 (17.2%) carried 2, and 5 (1.72%) carried 3 *dfrA* genes (**Fig 4c**). The diversity of *dfrA* genes highlights the adaptive potential of *P. mirabilis* to acquire and maintain resistance determinants. Resistance to sulfonamides (SMX) was similarly predicted in 286 (28%) genomes, with *sul1* detected in 180, *sul2* in 187, and 81 genomes harboring both *sul1* and *sul2* genes (**Fig 4d, Table S6**). Importantly, 223 of the 363 genomes encoding resistance genes for either TMP or SMX had resistance determinants for both (**Fig 4e**). Since TMP and SMX are often used in combination, the co-occurrence of these genes would likely complicate efficacy of combined treatment. Additionally, we detected 185 genomes (18%) with Extended Spectrum β-Lactamase (ESBL)-associated genes. Of these, 148 harbored the *blaTEM-1* gene, expected to confer resistance to ampicillin (**Table 3**).

Finally, the number of resistance subclasses detected per *P. mirabilis* genome ranged from 1 to 24, with an average of 4.61 AMR subclasses per isolate. A total of 71 genomes (7%) exhibited resistance to only a single antibiotic subclass, most commonly tetracycline. In contrast, 931 (93%) had resistance to two or more subclasses, and a substantial subset of 246 genomes (25%) harbored more than six antibiotic subclasses, indicating a high prevalence of multidrug resistance in this sample set. 176 genomes (18%) had at least one resistance determinant from all five of the above resistance gene groups, while 56 (6%) had combinations of tetracycline, chloramphenicol, and β-lactam resistance genes. Additionally, phylogenetic analyses (**Fig 1a**) determined that certain clusters consistently encoded resistance genes to nearly all antibiotic classes shown, such as genomes belonging to ST135, while others exhibited a complete absence of resistance genes other than tetracycline, independent of study source.

### Antibiotic resistance subclasses are lineage-associated but not strictly determined by ST

To evaluate the relationship between ST and AMR burden in *P. mirabilis*, we quantified the number of predicted AMR subclasses per genome. Logistic regression confirmed that MLST was not a significant driver of high AMR carriage overall (OR 0.0997, 95% Cl 0.992-1.002, p=0.304). However, striking lineage-specific enrichment was observed with stratification by resistance burden. Hence, to enable robust comparisons, we focused on a subset of 422 genomes representing the 16 STs with ≥10 genomes each. This accounted for 41% of the dataset (**Table S7, S8**). Within this subset, the mean AMR subclass count was 4.4 per genome, with most genomes carrying modest resistomes (80^th^ percentile ∼6 subclasses).

Genomes encoding resistances to ≥6 distinct subclasses were enriched in 5 STs (121/142 [85%]): ST135 (37/37 genomes [100%]), ST185 (33/47 [80%]), ST279 (10/12 [83%]), ST97 (25/36 [69%]), and ST145 (16/32 [50%]) (Pearson χ² p<0.001). Among these 142 genomes, 41 (29%) carried resistance genes to ≥12 subclasses, 33 of which (80%) belonged to ST135. In contrast, ST145, ST93, ST33, and ST202 typically carried 2-3 resistance subclasses with some outliers, highlighting inter lineage heterogeneity (**Table S8**). We acknowledge that sampling bias is possible as many surveillance studies only pick up on hospitalized or severe infections, but our dataset also includes general isolate submission from the CDC surveillance study, and we are most interested to identify clinically relevant strains.

ST135 emerged as an exceptionally high-risk clone; 95% of ST135 isolates (35/37) harbored ≥16 AMR genes, and 100% carried resistance to ≥6 subclasses. We next examined whether geographic origin confounded this association and created a bias. Within the 422-genome subset, 325 genomes had associated location data (US states/cities) and 167 were from St. Louis. Although a slightly higher proportion of St. Louis isolates exhibited high AMR carriage (≥10 subclasses) compared to other locations (3.4% vs. 0.9%; Pearson χ² p=0.037), isolation source was not an independent predictor in multivariable logistic regression controlling for MLST (OR 1.31, 95% CI 0.88–1.97, p=0.185). The strong enrichment of high-level AMR in ST135 regardless of geographical location points to lineage-intrinsic genetic factors as the primary drivers.

### The distribution of resistance gene burden is consistent across the dataset

To contextualize our 26 clinical isolates and HI4320 within the broader *P. mirabilis* population, we compare their AMR gene content against the 1,001 NCBI genomes. In total, resistance genes representing 14 different antibiotic subclasses were identified in the clinical isolates compared to 24 from the NCBI dataset. The 26 clinical isolates and Reference strain HI4320 (total 27) exhibited substantial variation in resistance gene burden, ranging from 1 gene (minimum, observed in ST765 and ST591 isolates carrying only a tetracycline resistance determinant) to 15 genes (maximum, in ST135 isolate H115-00, H115-29, and H127-00), with an average of 3.8 resistance genes per isolate (**Table 3 and Table S9**).

Despite these differences in dataset size and diversity, several resistance traits were highly conserved across both datasets. Tetracycline resistance approached fixation; *tetJ* was present in all isolates predicted to be tetracycline resistant from both datasets, and co-occurrence of *tetJ* with *tetA and tetC* (11%, 3/26 clinical) was also observed. Similarly, chloramphenicol resistance genes demonstrated strong cross-reservoir conservation, with *catA* (type-4) detected in 89% (23/26) of clinical isolates compared to 95% of NCBI genomes, and 26% of clinical isolates (6/23) co-harbored *catA1* while 11% (3/26) harbored no chloramphenicol resistance genes (**Table S9**). ST135 strains also exhibited a rare combination of *floR* + *catB3*.

Fewer of the clinical isolates had resistance genes for cephalosporins or quaternary ammonium in comparison to the NCBI dataset (**Table 3**). However, clinical isolates showed enrichment of resistance genes under UTI treatment pressure including TMP-SMX (22%, *sul1/sul2*; 37%, *dfr*) and aminoglycosides (streptomycin (37%, most common: *aadA1*), streptothricin (37%, *sat2*), and kanamycin (26%, *aph(3’)-Ia*) (**Table 3**). In terms of extended β-lactam ecology, clinical isolates predominantly carried *blaTEM-1* (18%) while NCBI genomes exhibited broader cephalosporin resistance genes (**Table 3, Table S6**). Collectively, these comparisons reveal a conserved genetic backbone of core resistance determinants (tetracycline, chloramphenicol) across *P. mirabilis* populations, upon which lineage-specific variation is superimposed.

### The mobile genetic element (MGE) landscape of *P. mirabilis* reveals an IS26-mediated major gene cluster in ST135

To determine if MGE’s are associated with AMR gene dissemination in *P. mirabilis*, we characterized 22 complete, polished reference quality genomes from the clinical cohort. We carried out MGE analysis only using only complete, polished genomes for high accuracy. MGE was not done for short read assembled genomes of the NCBI database since incomplete genomes it is not feasible to get an accurate picture from segmented genomes. Among the 22 isolates, a median of 22 IS elements per genome was detected (mean=23.68; range 16-36) that were distributed across 17 IS families (**Fig 5a, Table S7**). The most abundant family was IS200/IS605 (48.5%), followed by IS3 (16.31%), and ISNCY (7.2%). The three isolates with the highest total IS count (36 each) were ST135 (H115-00, H115-29, and H127-00).

**Fig 5.**
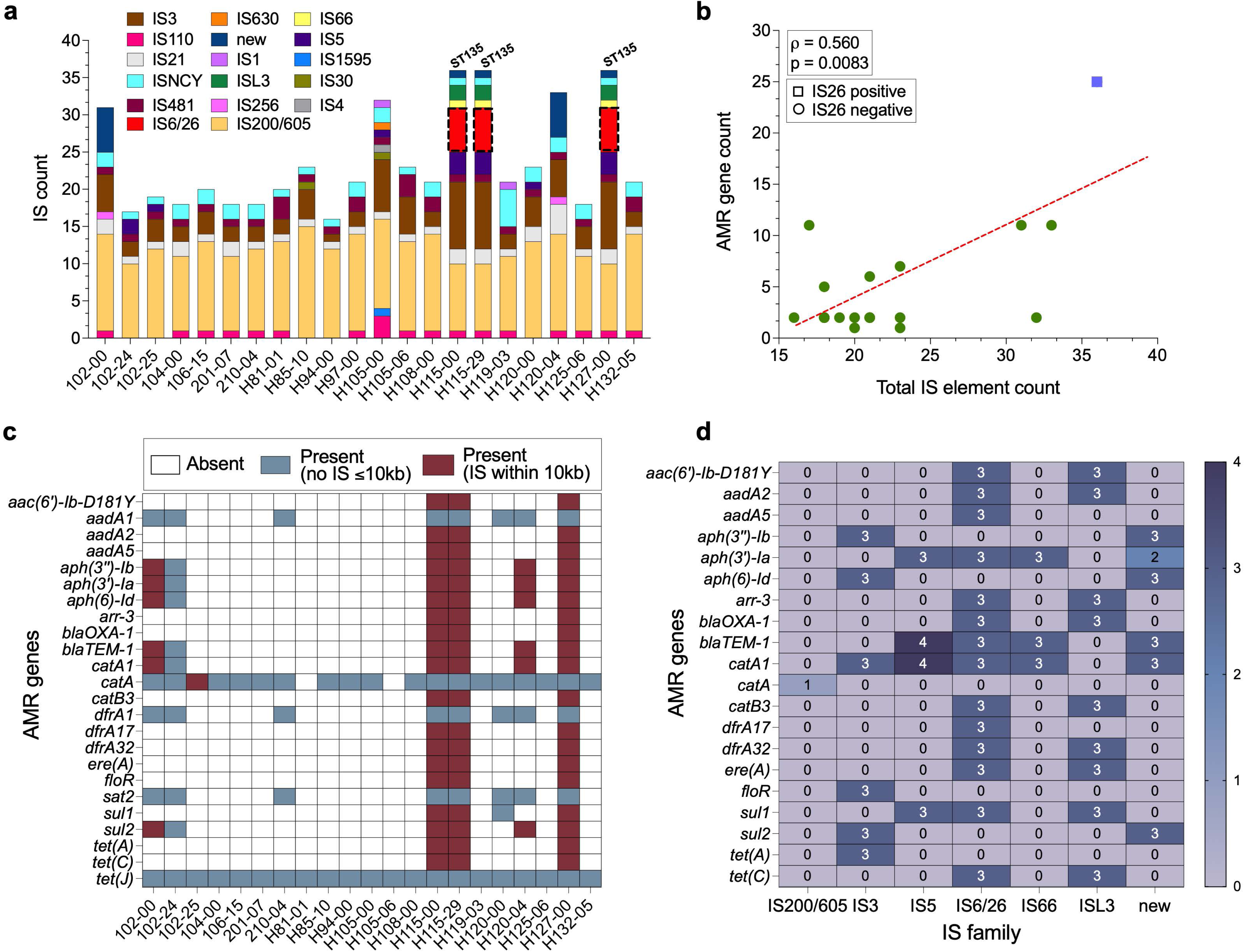
IS Element landscape overview of clinical isolates. **(5a)** Stacked bar chart of IS family distribution across STs. **(5b)** Scatter plot of total IS count vs. AMR gene count with Spearman correlation line (ρ=0.560, p=0.008). **(5c)** Proximity heatmap showing IS26 distance to each AMR gene in all isolates. **(5d)** Co-occurrence of IS family-AMR genes in isolates with IS within 10kb of AMR genes.

The high AMR gene count observed in the three ST135 isolates is positively co-related with IS element count (**Fig. 5b**). A large AMR gene cluster comprising 14 genes (*aadA2–ere(A)–dfrA32–sul1–arr-3–catB3–blaOXA-1–aac(6’)-Ib-D181Y–tet(C)–sul2–aph(3’’)-Ib–aph(6)-Id–tet(A)–floR)* was associated with flanking IS6 (also known as IS26) (**Fig. 5c, 5d**). This cluster spanned ∼72.6 kb of contiguous AMR gene content within an IS-bounded region of ∼180 kb. IS26 elements (3 copies, **Fig 5d**) were positioned within and flanking this cluster, with the closest IS26–AMR gene distances of 88–956 bp for six clinically important resistance genes including *aac(6’)-Ib-D181Y* (92 bp), *aph(3’)-Ia* (88 bp), and *sul1* (112 bp). This organization closely matches the *PmGRI1* genomic resistance island previously characterized in *P. mirabilis* Cluster-1 strains [64]. In contrast, non-ST135 isolates with moderate to low AMR burdens (e.g., ST185 with 11 genes, ST145 with 5–11 genes) lacked IS26 entirely (**Fig 5a**). IS elements flanking AMR genes in these isolates belonged primarily to the unclassified “new” family, IS3, and IS5 (**Fig 5d**).

A conserved Tn7-associated gene cassette (*aadA1-sat2-dfrA1*) was identified in 9/22 isolates (40.9%), three of which were ST135 (**Table S10**). This cassette displayed an identical 1,938 bp span and gene order in all positive isolates regardless of phylogenetic lineage, confirming the clonal stability of this segment. Interestingly, the three ST135 isolates also harbored a second AMR gene cluster (*catA1-blaTEM-1-aph(3’)-Ia-dfrA17-aadA5-sul1*) spanning ∼14.3 kb with identical gene content and order between genomes.

### The MGE landscape revealed Type IV secretion-system-type integrative and conjugative elements (ICE) in *P. mirabilis* genomes

Genome mining using ICEberg 3.0 [62] identified 1-2 Type IV secretion system (T4SS)-type ICEs in the 22 complete, polished chromosomes of *P. mirabilis*. The GC content of these elements ranged from 44.86%-46.19%, they integrated near the tRNA-Phe gene, flanked by identical 52-bp direct repeats (attL/attR), and carried a mobilization protein family H (MOBH) relaxase and a type G mating pair formation system. ST135 isolates (H115-00, H115-29, H127-00) carried one identical ICE harboring 13 AMR genes along with a copy of the IS26-flanked cluster of 5 genes (*aacA4-blaTEM-catA1-emrE-folP1*) (**Fig 6a**). In contrast, ST185 isolates (102-00, H120-04) had 2 regions with T4SS-type ICEs. Annotation of the ICE identified a range of virulence, housekeeping, and metabolic genes (e.g., permease, PikA5); **Fig. 6b**). The other isolates carried only one ICE, harboring similar virulence-associated genes as the region 1 ICE in ST185 (**Fig. 6c**).

**Fig 6.**
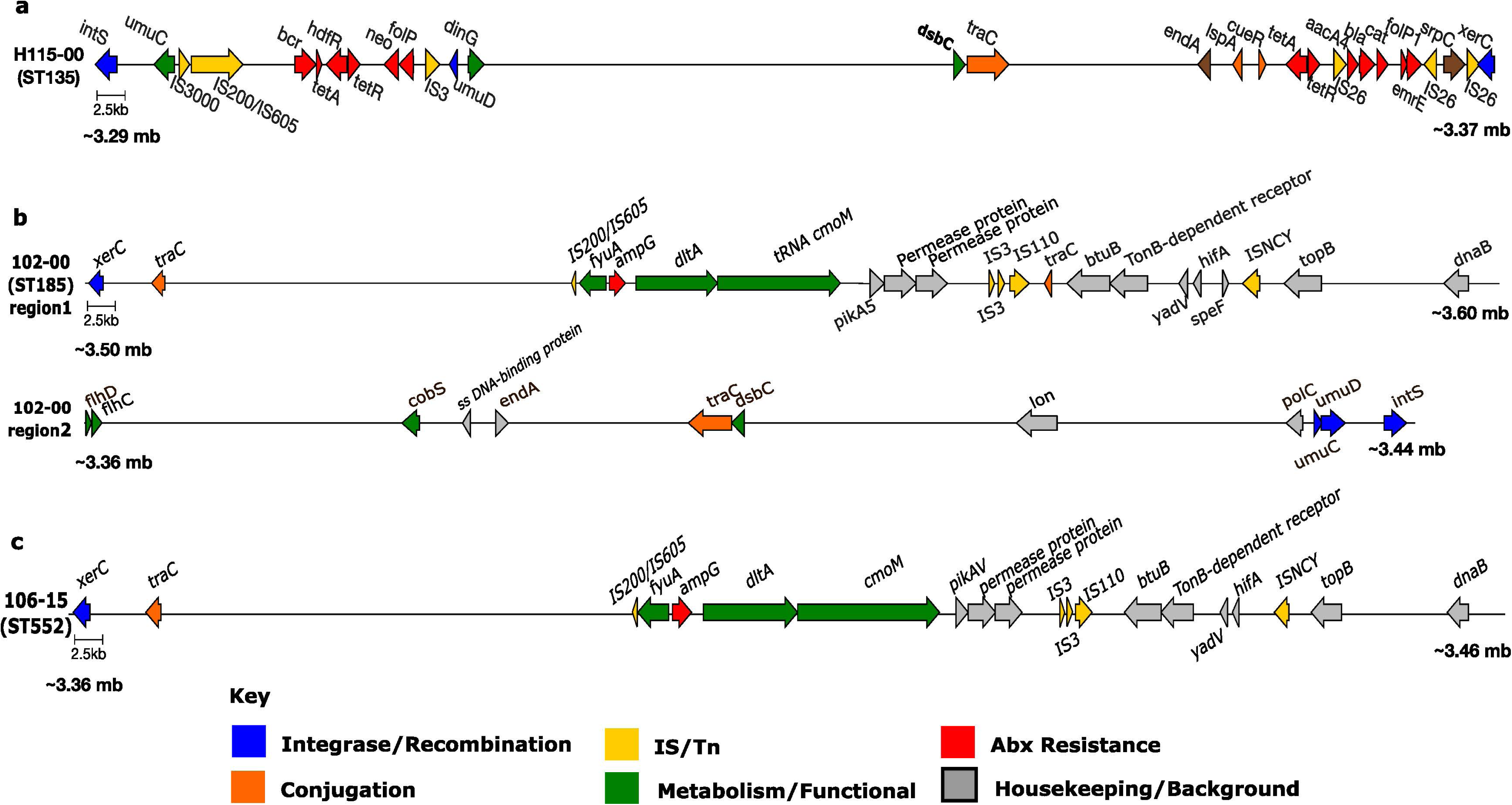
Genetic organization of the T4SS integrative and conjugative elements (ICEs) identified in *Proteus mirabilis*. (**6a**) Genetic structure of the ICE identified in strain H115-00. The element includes the integrase gene *intS* and regulatory gene *umuC*, followed by insertion sequence elements (IS3000, IS8 family) and multiple accessory genes. A cluster of antibiotic resistance genes is present, including genes such as *bcr, tetR*, *tetB, dfrA, aac, aph*, and *erm* family members, interspersed with transposition– and recombination-related genes (*tnpA, xerC).* **(6b)** 2 T4SS-ICEs detected in strain 102-00. Genes are shown as arrows indicating the direction of transcription. The element contains recombination genes *recC* and *uvrC*, followed by multiple accessory and housekeeping genes. Conjugation-associated genes including *tra* components are present toward the right side of the element together with additional metabolic genes. **(6c)** T4SS-ICE detected in strain 106-00. This is a representative for all the non-ST135 and ST145 in our dataset. The ICE is similar to the first ICE observed in ST145 in 6b.

### Prophages and plasmids have a minimal contribution to AMR gene carriage in *P. mirabilis*

PHASTER analysis of the 22 clinical isolate genomes identified 189 prophage regions (mean: 8.2/genome), including 64 intact (33.8%), 20 questionable (10.5%), and 104 incomplete (55%) (**Table S10**). Coordinate-overlap analysis revealed 14 AMR gene instances within prophage regions across 3/11 isolates (27%), none of which were ST135. These AMR genes in these prophage regions included 10 genes spanning six resistance classes (aminoglycosides, β-lactams, chloramphenicol, sulfonamides, trimethoprim, and disinfectants).

PlasmidFinder detected replicons in only 2/26 isolates (6.7%): IncN4 (conjugative) plus Col3M and ColE10 in H94-00 (ST696), and a single ColE10 in H81-01 (ST765) (**Table S10**). The only confirmed plasmid-borne AMR gene across all 30 isolates was *qnrD1* (quinolone resistance) on the Col3M plasmid in H94-00 (ST696). This indicates that the overwhelming majority of AMR in this *P. mirabilis* collection are chromosomally encoded.

### Hierarchical MGE overlaps reveal a “Russian doll” architecture driving multidrug resistance in ST135

Systematic analysis of MGE overlaps within the 22 clinical isolates revealed a nested hierarchical architecture most pronounced in ST135 isolates. Class 1 integrons with their 3’-CS (*qacE*Δ*1-sul1*) are embedded near IS26 sites. Each integron carries distinct aminoglycoside and trimethoprim resistance cassettes (*aadA, dfrA*). These modules are themselves clustered within *PmGRI1-*like genomic islands containing IS elements. In ST135 isolates, IS26 serves as the structural organizer of *PmGRI1*: Region 1 contains 6 IS elements (3× IS26, 2× ISL3, 1× IS3) interspersed among 16 AMR genes (**Fig 7a**); Region 2 contains 8 IS elements (3× IS26, 3× IS5, IS66, IS200/IS605) among 10 AMR genes (**Fig 7b**). IS26 may act as a boundary element for sub-modules within *PmGRI1*, enabling modular rearrangement and stepwise accumulation of resistance. In the same arrangement, Prophage region 10 (35.1 kb, intact, score 130, with *att* sites) simultaneously overlaps region 2. This represents a four-way overlap of all element types within a single 35 kb region, nested within *PmGRI1*.

**Fig 7:**
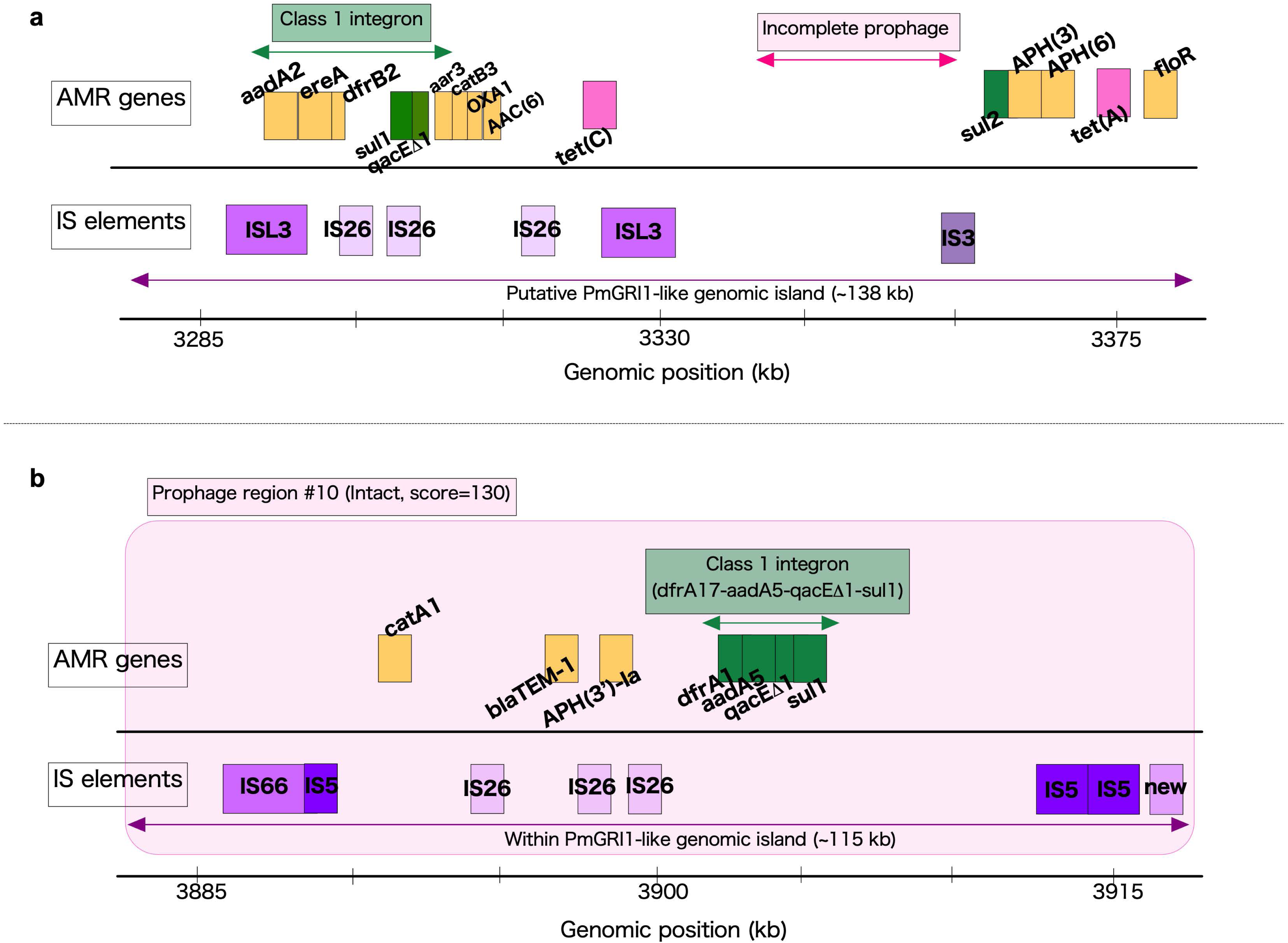
Comprehensive map of the *Proteus mirabilis* Genomic Resistance Island 1 (PmGRI1)-like region in H115-00 (ST135): **(7a)** Putative PmGRI1 region 1 (3.29 Mb-3.37 Mb), and **(7b)** Putative PmGRI1 region 2 (3.885 Mb – 4.0 Mb). PmGRI1 is a chromosomally integrated, modular resistance island identified by tyrosine-family site-specific recombinase/integrase (∼394 aa), insertion at the 3’ end of tRNA-Sec, and 20bp direct repeats flanking the element.

### Experimental AMR phenotyping of *P. mirabilis* clinical isolates reveals variable genotype-phenotype concordance

AMRFinderPlus reports gene presence/absence and point mutation but does not infer phenotypic resistance due to factors including gene expression and regulation. Thus, many of the resistance genes detected by established computational databases may not be relevant for clinical management or antimicrobial surveillance. To determine concordance between *in-silico* AMR predication and phenotypic resistance in *P. mirabilis*, we determined the AMR of all 27 *P. mirabilis* clinical isolates by standardized Antimicrobial Susceptibility Tests (AST). We tested growth kinetics of strains with specific resistance genes at 15-minute intervals over a 20-hour time course to evaluate the minimum inhibitory concentrations as well as more nuanced dose-dependent effects on the growth rates and doubling times of the *P. mirabilis* strains (**Fig 8 and Fig 9**). Antibiotic concentrations were selected based on the Clinical and Laboratory Standards Institute (CLSI) guidelines for *Enterobacterales* and were designed to span the thresholds for sensitive (lowest) to resistant (highest). The genotypic and phenotypic resistance profiles are summarized in **Table S9 and S11.**

**Fig 8:**
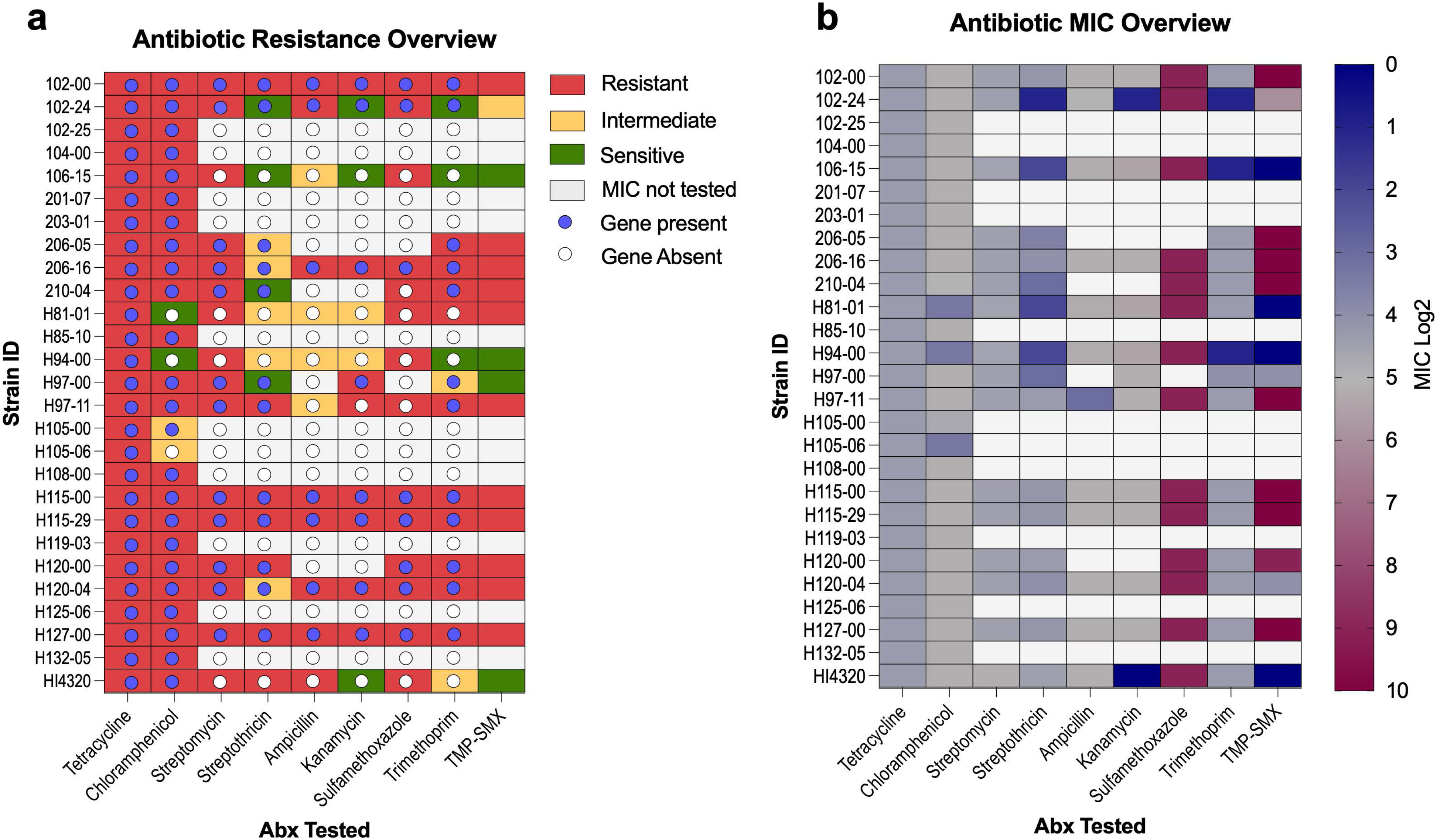
Overview of antibiotic resistance and MIC. (**7a**) Overview of antibiotic resistance phenotype found in the clinical CAUTI isolates. Color coding denotes resistance status: red = resistant, yellow = intermediate resistance, and green = susceptible strains. The presence or absence of resistance-associated genes is indicated by filled and empty circles, respectively. (**7b**) Heatmap representing the MIC for each antibiotic tested across clinical *P. mirabilis* CAUTI isolates. Color intensity corresponds to MIC values, indicating variation in resistance levels.

**Fig 9:**
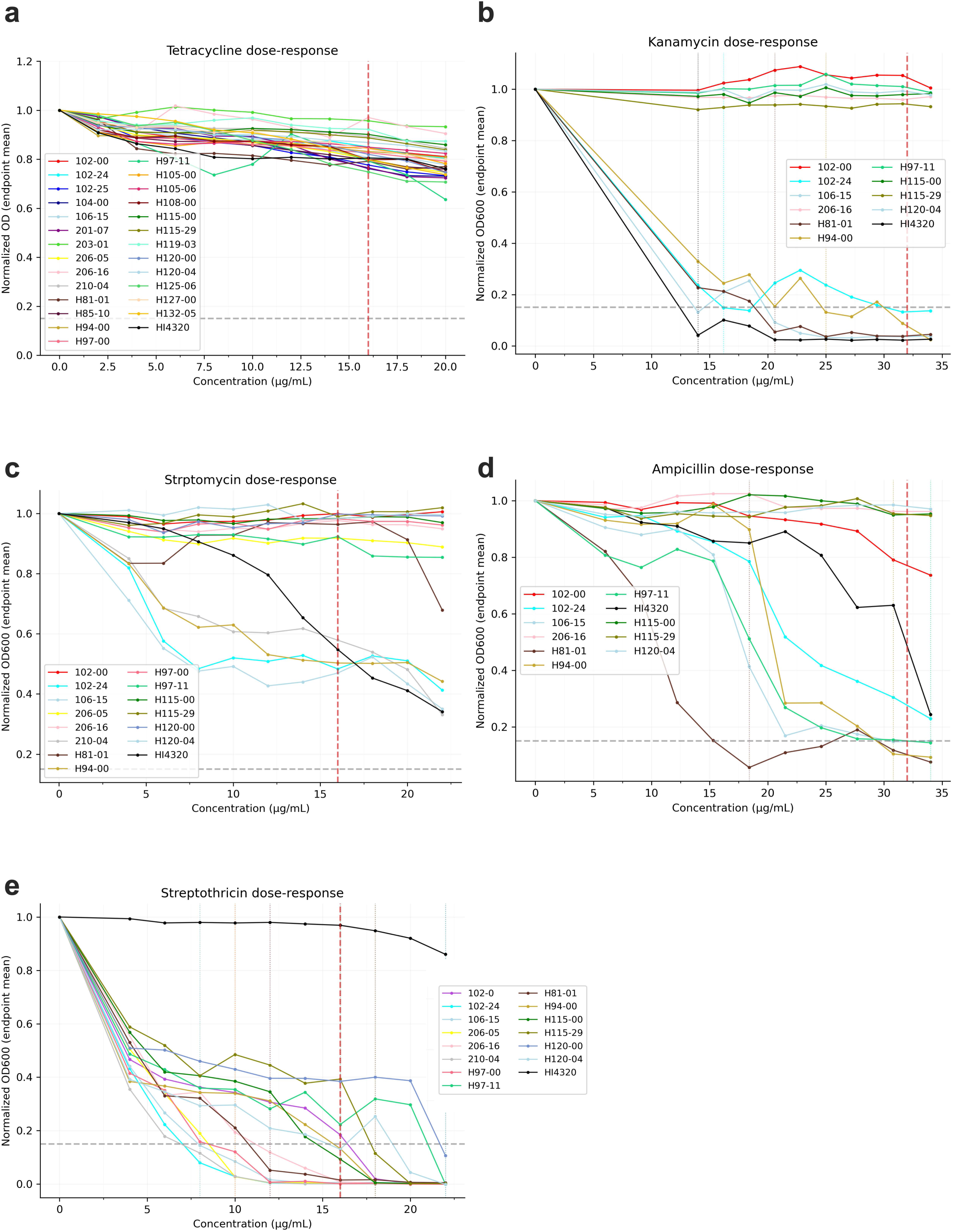
Dose response curves and MIC. Mean normalized OD600 values at experimental endpoint (20 hours) are plotted against antibiotic concentrations in ug/ml. OD is normalized to a no drug control set to 1.0 (maximum growth). The red vertical line indicates the resistance breakpoint (CLSI) for *Enterobacterales.* The MIC was determined as the lowest antibiotic concentration for which growth was restricted below a threshold of a normalized OD of 0.15 (grey horizontal dashed line). (**8a**) Tetracycline (MIC range tested: 2-20 ug/ml), (**8b**) Kanamycin (MIC range tested: 6-34 ug/ml), (**8c**) Streptomycin (MIC range tested: 4-22 ug/ml), (**8d**) Ampicillin (MIC range tested: 6-34 ug/ml), (**8e**) Streptothricin (MIC range tested: 4-22 ug/ml).

#### Tetracycline, Kanamycin, Streptomycin, Streptothricin, and Ampicillin

For tetracycline, 100% of clinical isolates (26/26) exhibited resistance with MIC values >20 μg/mL, which is consistent with the presence of *tetJ*. Growth kinetics demonstrated uniformly robust growth across all concentrations (**Fig 9a**). Thus, a perfect genotype-phenotype concordance was observed. Carriage of any additional MGE-acquired *tet* genes (*tetA* or *tetC*) are phenotypically redundant.

Kanamycin resistance genes (*aac(6’)-Ib-cr5, aph(3’)-Ia*) were detected in 7/26 clinical isolates and genotype-phenotype concordance was observed. All 7 isolates encoding the resistance genes had MIC >34 ug/mL, while 106-15, H81-01, H94-04, and HI4320 were predicted to lack all kanamycin resistance determinants and were susceptible, but some exhibited MICs ranging from 14 – 20.5 ug/ml (**Fig 9b**).

Different combinations of streptomycin resistance genes (*aadA1, aph(6)-Id, aph(3”)-lb, aadA5, aadA2*) were observed in 11/26 clinical isolates (**Fig 8a, Table S10**), and all provided resistance to streptomycin when tested in our AST assay (**Fig 9c**). Unexpectedly, resistance was also observed in 102-24, 106-15, H81-01, H94-00, and reference strain HI4320 which lacks canonical streptomycin resistance genes. However, these strains all exhibited a dose-dependent decrease in growth when incubated with streptomycin. It is also notable that clinical isolate 210-04 (1 predicted gene, *aadA1*) exhibited a similar dose-dependent decrease in growth even though other strains (206-05, H97-00, and H97-11) with a singular *aadA1* gene with 100% amino acid sequence similarity to that of 210-04 did not show dose dependency. It is possible that regulation of *aadA1* differs in 210-04, leading to a less efficient or inducible mechanism of resistance compared to the other isolates. The presence of multiple, stacked aminoglycoside resistance genes on MGEs like *PmGRI1* in H115-00 (e.g., *aadA1, aph(6)-Id, aph(3’’)-lb)* can confer broader and potentially higher-level resistance compared to strains carrying only a single gene, explaining quantitative differences in MICs beyond simple gene presence.

An ampicillin resistance gene (*blaTEM*) was identified in 6/26 clinical isolates and all exhibited MICs >34 ug/mL. The resistance phenotype was also observed in HI4320, which lacks any known β-lactamase genes. However, strains that did not encode *blaTEM*, including HI4320, exhibited dose-dependent growth inhibition with increasing concentrations of ampicillin, despite displaying resistance at CLSI breakpoints (**Fig 9d**). The dose-dependent growth inhibition observed in some strains with *bla*TEM, and the low-level resistance in strains lacking it, may be due to IS-proximal regulatory effects influencing gene expression or the presence of alternative, non-canonical resistance mechanisms located within genomic islands.

Streptothricin resistance gene *sat2* was observed in 11/26 clinical isolates (**Fig 8a, Table S9**), yet the AST assay revealed a mix of intermediate to resistant phenotypes with MICs ranging from 8 to >22 ug/mL (CLSI breakpoints are ≤8 for susceptible and ≥16 for resistant). Since streptothricin resistance is conferred by *sat1, sat2*, and *sat4* genes, the presence of only one gene may confer only partial resistance with dose-dependency increasing with gene stacking. The genotypically-positive isolates 102-00, 206-16 H97-11, H115-00, H115-29, H120-00, and H120-04 exhibited dose-dependent decreases in growth with increasing streptothricin concentrations. However, other genotypically positive strains 206-05 and 210-04 were sensitive (**Fig 9e**). HI4320 exhibited robust growth regardless of streptothricin concentration, yet this strain does not encode any known streptothricin resistance genes (**Fig 9e).**

#### Sulfonamide and Trimethoprim

Sulfonamide resistance genes (*sul1* and *sul2)* were both observed in the 26 clinical isolates, with *sul1* present in 3 isolates (H115-0, H115-05, H120-00), *sul2* in 5 isolates (102-00, 206-16, H120-04, H115-00, H115-05), and both genes co-occurring in 3 isolates (H115-00, H115-05. H127-00) (**Table S9**). AST with sulfamethoxazole revealed high level resistance (MIC >514 ug/ml) in all isolates, irrespective of *sul* gene carriage (single or double), indicating that the presence of an additional *sul* gene did not further elevate resistance. Unexpectedly, even strains that did not encode *sul* (102-24, 106-15, 210-04, H81-01, H94-00, H97-11, and HI4320) had MICs >514 ug/ml, but partial growth inhibition was clearly present even at the lowest tested concentration (250 ug/ml, **Fig. 10a**). The uniform MIC >514 µg/mL across all strains implies clinical resistance per CLSI guidelines, as it far exceeds the susceptible breakpoint (typically ≤38 µg/mL). However, the partial growth inhibition of strains that do not encode *sul* highlights the complexity of interpreting MICs near the resistance threshold. These data suggest that, while canonical *sul*-mediated resistance mechanisms were absent in these isolates, IS elements located near AMR genes or regulatory regions may contribute to the observed resistance (above threshold) phenotype.

**Fig 10:**
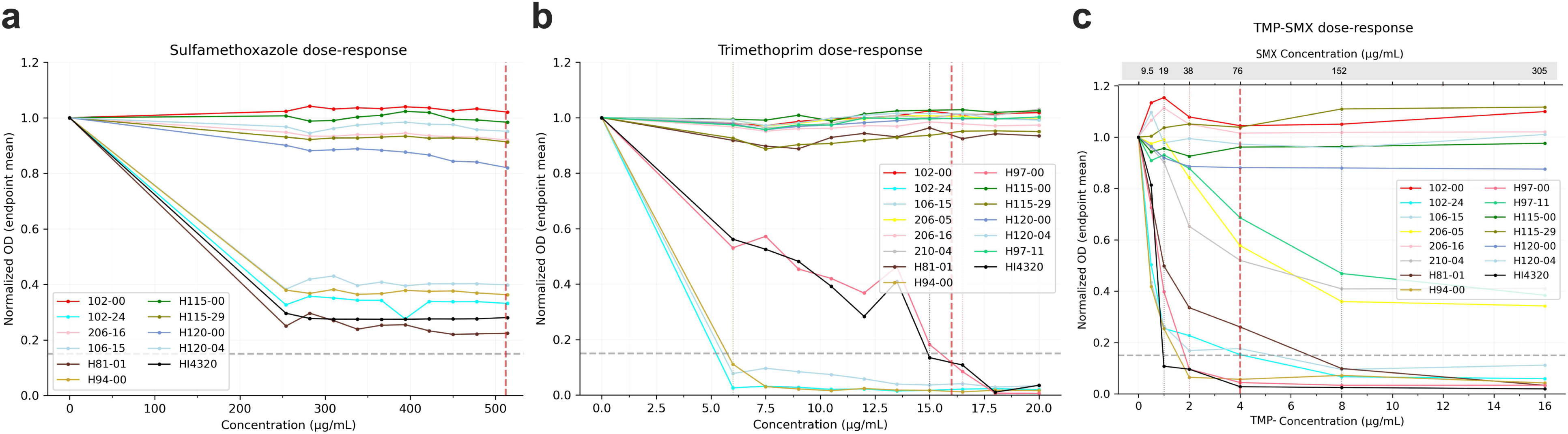
Dose response curves and MIC. Mean normalized OD600 values at experimental endpoint (20 hours) are plotted against antibiotic concentrations in ug/ml. OD is normalized to a no drug control set to 1.0 (maximum growth). The red vertical line indicates the resistance breakpoint (CLSI) for *Enterobacterales.* The MIC was determined as the lowest antibiotic concentration for which growth was restricted below a threshold of a normalized OD of 0.15 (grey horizontal dashed line). (**9a**) Sulfamethoxazole (MIC range tested: 250-512 ug/ml), (**9b**) Trimethoprim (MIC range tested: 4-22 ug/ml), (**9c**) TMP-SMX (MIC range tested: 0.5/9.5-16/305 ug/ml).

Trimethoprim resistance genes were also detected in 10/26 clinical isolates, with MICs of >15 µg/mL (exceeding clinical breakpoints) regardless of which resistance gene (*dfrA1*, *dfrA17*, or *dfrA32*) was present **(Fig 10b)**. However, one isolate (H97-00) harboring *dfrA1* exhibited dose-dependent growth inhibition, contrasting with other strains harboring *dfrA1* with 100% amino acid similarity that showed no dose-dependency. This implies strain-specific phenotypic variability (e.g., differential gene expression, compensatory mutations) despite shared genetic determinants. The *dfrA*-negative reference strain H81-01 also displayed full resistance, while strain HI4320 that did not carry the *dfrA* gene or gene mutation displayed intermediate resistance and mirrored the dose-dependent growth inhibition of strain H97-00. These observations support the existence of non-*dfrA* resistance mechanisms in HI4320 and H81-01, such as altered folate metabolism or reduced drug uptake. The discordance between genetic markers and phenotypic outcomes also underscores a key challenge: genetic testing alone cannot always predict antibiotic response. However, *dfr* negative strains 102-24, 106-15, and H94-00 were sensitive to trimethoprim, in concordance with the genotype.

Since the genotype-negative strains were all phenotypically resistant to sulfamethoxazole (SMX), albeit with reduced growth rate, we further tested the impact of combinatory treatment (trimethoprim-sulfamethoxazole, TMP-SMX) on *P. mirabilis* growth. This is an important parameter since, historically, TMP-SMX was the first-line empirical treatment for acute uncomplicated UTIs. In the strains exhibiting 100% genotype-phenotype concordance for both SMX and TMP resistance (i.e. true resistant genotypes correlated with resistant phenotypes), all isolates (102-00, 206-16, H115-00, H115-05, H120-00, and H120-04), exhibited complete resistance to the combinatory drug without any dose-dependent decreases in growth (**Fig 10c**). In contrast, strains such as 206-05, 210-04, and H97-11 that were genotypically SMX susceptible TMP resistant but phenotypically resistant to both antibiotics when used alone exhibited a dose-dependent resistance phenotype when treated with TMP-SMX. Additionally, strains that were genotypically predicted to be sensitive to both antibiotics but were phenotypically resistant demonstrated intermediate to resistant phenotypes for combination treatment: H81-01 displayed an MIC of 8/152 ug/mL (resistant), while 102-24 displayed an MIC >2/38 ug/mL (intermediate). This is clinically concerning as growth was observed at breakpoint concentrations. Other discordant strains that had lower growth yields under single SMX exposure (106-15, H94-00, H97-00, HI4320) were found to be sensitive to the TMP-SMX combinations (**Fig 10c**).

#### Chloramphenicol

Chloramphenicol resistance displayed genotype-phenotype concordance, with all isolates possessing a *catA* gene exhibiting resistance and those lacking *cat* genes exhibiting susceptibility (**Fig 11**). MIC values ranged from 10-14 ug/ml in isolates lacking a *cat* gene but were >26 ug/ml in isolates with *catA* alone and >34 ug/ml in isolates harboring both *catA* and *catA1* (**Table S9**). Interestingly, many isolates with a single *catA* gene (16/17) exhibited dose-dependent growth inhibition (**Fig 11**), while isolates possessing both *catA* and *catA1* demonstrated hyper-resistance (hR) with no dose dependency (**Fig 11**). The exception was HI4320, which only encodes *catA* but exhibited robust growth at all but the highest three concentrations of chloramphenicol. No strains were predicted to have *catA1* alone, so we were not able to determine whether *catA1* provides a greater level of resistance than *catA,* but our data suggests that the presence of two *catA* genes enhances functional resistance.

**Fig 11:**
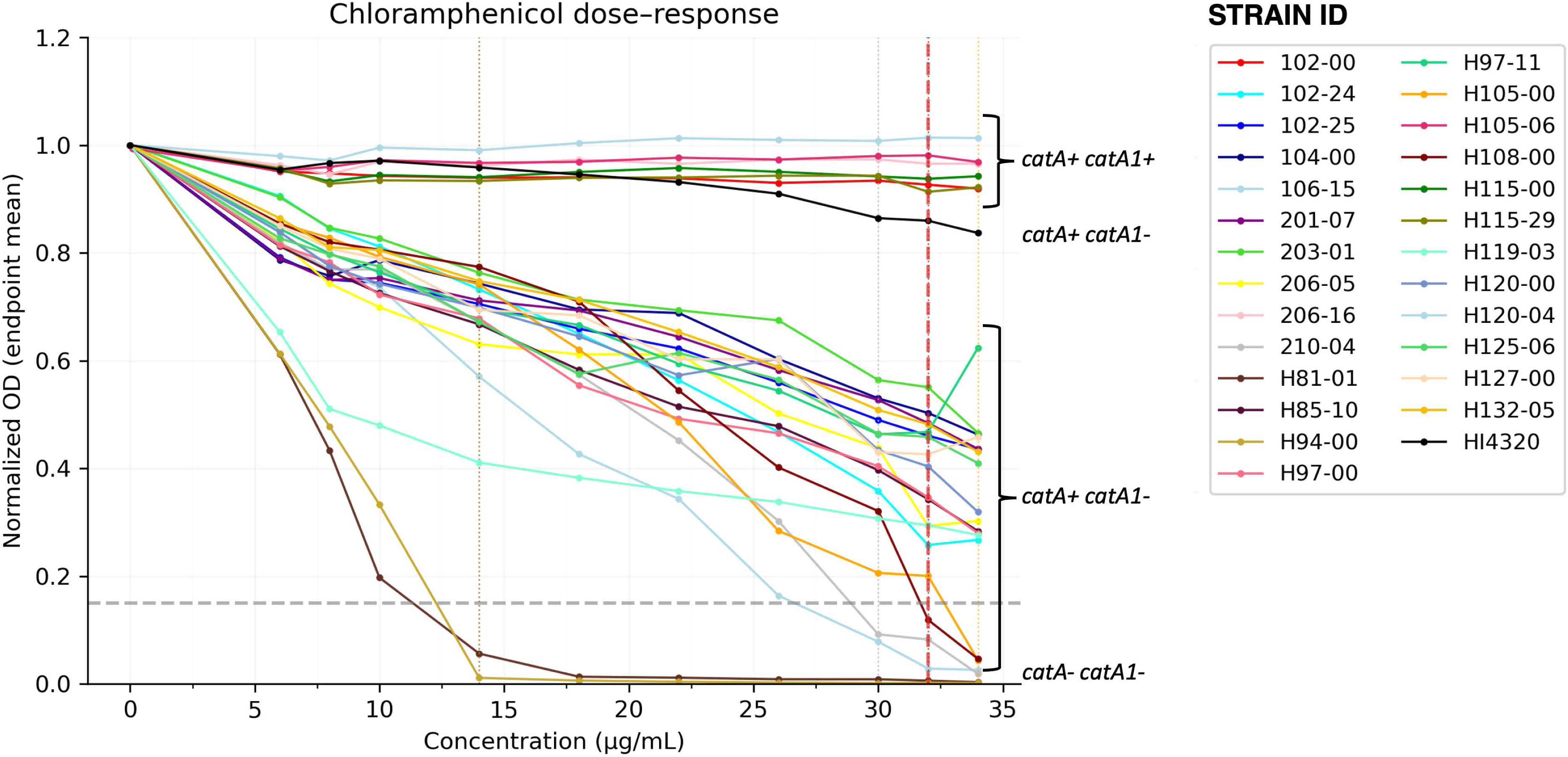
Dose response curves and MIC of Chloramphenicol. Mean normalized OD600 values at experimental endpoint (20 hours) are plotted against antibiotic concentrations in ug/ml. OD is normalized to a no drug control set to 1.0 (maximum growth). The red vertical line indicates the resistance breakpoint (CLSI) for *Enterobacterales.* The MIC was determined as the lowest antibiotic concentration for which growth was restricted below a threshold of a normalized OD of 0.15 (grey horizontal dashed line). Chloramphenicol concentrations tested ranged from 6-34 ug/ml.

We next compared the sequences of the *catA* and *catA1* genes in our *P. mirabilis* clinical isolates that harbored both genes as *catA1* has not previously been reported in *P. mirabilis*. The two genes share 76% amino acid sequence identity (with representative strain 102-00; **Fig S2a**), and *catA* from strain 102-00 showed 100% identity to the *catA* gene in reference strain *P. mirabilis* HI4320 (**Fig S2b**). The nucleotide and amino acid sequences of *catA1* from our isolates share 100% amino acid identity with *catA1* from *E. coli* and ∼98% amino acid identity with *catA1* from *Photobacterium damselae* and *Shigella flexneri* (**Fig S2c)**.

Even though we had a limited sample size, within-lineage discordance was observed in ST145 and can be attributed to differential MGE acquisition. 102-24 carries 11 AMR genes (including IS5-flanked accessory cluster with *aph(3’’)-Ib, aph(3’)-Ia*, *aph(6)-Id, blaTEM-1, catA1, sul2*) while 210-04 carries only 5 genes. The 6 additional genes in 102-24 are MGE-acquired (flanked by IS5 elements), conferring predicted resistance to 3+ additional drug classes. This means two isolates of the same sequence type would have different AST profiles, a difference that is possibly explained by differential MGE acquisition.

## DISCUSSION

CAUTIs caused by *P. mirabilis* remain clinically challenging due to crystalline biofilm formation, recurrent infections, and persistence despite antibiotic treatment and catheter changes [15]. This comprehensive genomic analysis of *P. mirabilis* from human urinary sources expands our understanding of *P. mirabilis* genetic diversity, AMR profiles, and the accessory genome. We analyzed 1,027 publicly available and clinical isolates and determined that urinary *P. mirabilis* isolates possess an open pangenome with more than 41,121 gene families and a core genome of 1,937 genes. Open pangenomes are characteristics of sympatric species with a high capacity of horizontal gene transfer (HGT) mediated by MGEs, highlighting genomic plasticity. This may allow *P. mirabilis* to rapidly adapt to the urinary tract under antibiotic exposure, host immune responses, and competition in the polymicrobial catheterized environment [65–67]. This dynamic gain, loss, and exchange of MGEs with accessory genomes comprising ∼90% (37,918 genes) emphasizes the importance of flexible genomic elements in driving diversity in the urinary tract.

The presence of 213 defined STs and ∼118 previously undefined genomes and a star-like phylogeny further supports ongoing diversification. Almost half of the STs were represented by single isolates (46%), showcasing heterogeneity. Yet, most genomes were distributed across a small subset of STs (ST96, ST185, ST99, ST135, ST97), resulting in a skewed distribution. While MLST remains a widely used typing method due to its reproducibility and interpretability [51, 68], we want to highlight its limitations for epidemiological investigations [68, 69]. The substantial proportion of unidentified STs and limited sample size constrain ST-specific characterization in the entire 1,027 genomes but highlights the need for continuously updated typing schemes.

The AMR landscape was complex and dynamic, with variable genotype-phenotype concordance. Overall, 93% of the isolates were resistant to ≥2 antibiotic subclasses and 25% resistant to >6 subclasses posing a challenge for effective treatment of *P. mirabilis* UTI’s. Although MLST was not a significant predictor of high AMR carriage in the full dataset of 1,001 genomes, ST135 emerged as a high-risk clone is the subset of 422/1,001 with >10 genomes. 95% of ST135 genomes had ≥16 resistance genes and 100% had resistance to >6 antimicrobial subclasses (**Fig 1a**). This near-uniform high-level resistance within ST135 is attributed to the IS26-mediated MGEs that are rarely observed in other lineages. IS6/IS26-encoding genomes carry 3.2× more AMR genes (20.8 vs 6.5) and 2.8× more drug subclasses (13.9 vs 4.9), with a significant dose-response relationship as IS6 copy number increases. MLST ST135 had the highest IS diversity in our dataset (37/422 genomes + 3 clinical isolates), highest IS6/IS26 prevalence, and AMR gene burden with 100% MDR (>3 antibiotic subclass).

Geographic confounding was excluded by our analysis; although ST135 was predominantly isolated from St. Louis, isolation source alone was not an independent driver of high AMR carriage (p=0.051) pointing to lineage-intrinsic factors rather than regional antibiotic selection pressures. The absence of geographical associations, together with the hierarchical MGE architecture we identified (**Fig 7**) implicates lineage-specific MGE carriage as the mechanistic basis for extreme resistance phenotypes. This is known for other species as well—*S. aureus* USA300 (ST8/5) carry the mecA cassette and are seen globally [70–72]. The overlapping MGE architecture (H115-00; **Fig 7**) reveals a multi-level system for AMR propagation. Class 1 integrons capture individual resistance cassettes, IS26 organizes these into modular translocatable units, *PmGRI1* provides stable chromosomal integration, and prophages enable horizontal transfer of complete AMR modules. This nested hierarchy has clinical implications because redundant dissemination pathways can ensure robust AMR spread. Clustering of multiple resistance genes within shared elements means selection for any single antibiotic co-selects the entire MDR module, and chromosomal integration ensures stable inheritance while IS26 and phage mechanisms retain mobility. The exclusive association of this architecture with ST135, which carries significantly more AMR genes than IS26-negative lineages (mean 25.0 vs. 4.1, p=0.0019), implicates these overlapping MGEs as key drivers of this lineage’s epidemiological success.

A recent study by *Zhang et. al. (2026)* identified the same high-risk lineage which they assigned Cluster-1 genomes comprising 233 globally distributed isolates [64]. Our analysis confirmed that Cluster-1, which mostly carries ST135 (159/233), had significantly elevated AMR gene burden. The high AMR gene counts across isolates from different locations and hosts that was reported by *Zhang et. al.* further supports ST135 as a lineage of major clinical concern [64]. The T4SS ICE and IS26-mediated AMR gene clusters help to stack AMR genes within *PmGRI1*, enabling vertical transmission and potentially facilitating further resistance acquisition. Such lineage-specific AMR enrichment has critical implications for infection control and genomic surveillance, as clonal expansion of ST135 could rapidly disseminate multidrug resistance across healthcare networks like that or Methicillin resistant *S. aureus* (MRSA), Vancomycin resistance *E. faecalis* (VRE) [70, 72–78]. Prospective surveillance should therefore prioritize tracking this high-risk lineage, and future functional studies should dissect the precise MGEs responsible for its resistance burden and why is it limited to this specific lineage. In contrast, the broader distribution of common resistance subclasses across multiple STs reflects ongoing horizontal gene transfer. These findings highlight the critical need for genomic surveillance targeting high-risk lineages, as clonal expansion of ST135 could rapidly disseminate multidrug resistance across healthcare networks.

The overwhelming prevalence of tetracycline resistance genes (98.9%), particularly *tetJ*, aligns with intrinsic resistance of *P. mirabilis.* The dominance of *tetJ* and the frequent carriage of only a single *tet* gene (94.3%) suggest this determinant is often sufficient for the observed resistance phenotype. Tetracycline resistance also exemplifies how MGE-acquired genes can be phenotypically redundant: *tet*(J) alone confers complete resistance (MIC >20 µg/mL), yet some genomes such as the ST135 isolates carry three genes (*tet*(J), *tet*(A), *tet*(C)) acquired on the PmGRI1 genomic island via IS26. Thus, the additional MGE-borne genes inflate genotypic counts without meaningfully expanding the phenotype.

Chloramphenicol resistance genes (*catA*, 95%) were prevalent despite limited clinical use, possibly reflecting co-selection with other antibiotics or acquisition due to extensive use in agricultural and environmental settings [79, 80]. The co-occurrence of *catA* and *catA1* was detected in 19.2% of *cat*-positive isolates, with rare *floR* and *catB3* gene stacking. Strains with only a single *cat* gene exhibited dose-dependent growth inhibition, while additional *catA1* resulted in a hyper-resistant phenotype (no growth inhibition at all at any tested concentrations), enhancing the resistance phenotype. This observation suggests that the combined expression of multiple acetyltransferases, driven by IS26 elements positioned within 10 kb, quantitatively enhances resistance. Detection of *catA1* in *P. mirabilis* is a significant finding as this gene has not been well reported and characterized in *P. mirabilis* but match closely to related bacterial species [81, 82].

Additionally, we observed resistance to aminoglycosides (71%) with streptomycin resistance often co-occurring with other determinants of carbapenems, cephalosporins, fluoroquinolones and trimethoprim-sulfonamides (**Fig 1**). This persistence despite reduction in clinical use may reflect MGE mediated clonal spread of resistant lineages. Trimethoprim (TMP, *dfr*, 29%) and sulfonamide (SMX, *sul*, 28%) resistance determinants also co-occurred in 22% of genomes, threatening the efficacy of this combination therapy for *P. mirabilis* for UTI treatment. While ESBL-associated genes were less common (18.5% and primarily *blaTEM-1*), their presence is concerning for broader β-lactam resistance development. Resistance for quinolones (*qnr aac(6’)-Ib-cr, gyrA/parC* mutations) and the chloramphenicol/florfenicol (*cat, floR*) combination were rare in both datasets (∼7%) but noteworthy as potential molecular fingerprints of human driven selection linked to agriculture use and MGE gene transfer from *P. mirabilis* lineages found in animal sources [35, 64, 83–86].

Our comprehensive AMR analysis shows a complex genotype-to-phenotype landscape with both expected concordance and clinically critical discordance. Concordance for antibiotics like tetracycline was expected and kanamycin resistance reliably predicted by the presence of an *aph(3’)-la* gene, validated the robustness of our genomic approach for identifying known resistance determinants [75, 87, 88]. However, discordances provide compelling insights into AMR dissemination. For example, resistance to streptomycin and streptothricin could be due to novel acetyltransferases. Similarly, the ampicillin resistance in strains 102-24, 106-15, H94-00, H97-11, and HI4320 that lack a detectable β-lactamase gene may reflect permeability or efflux mechanisms, which is a hallmark of concentration-dependent antibiotics [89, 90], where higher concentrations or adjuvants can eventually overcome cellular defenses.

A critical finding was the discordance between genotypic SMX susceptibility and TMP-SMX phenotypic response (**Fig. 8**). Strains that lacked the canonical *sul* genes yet exhibited variable resistance phenotypes could possibly be explained by MGE-related mechanisms: 1) IS elements positioned ≤100 bp from AMR genes in seven isolates may drive upregulation via outward-facing promoters; 2) ST135 carries three *dfr* genes (*dfrA1*, *dfrA17*, *dfrA32*) accumulated via IS26 on *PmGRI1*, while some isolates carry only *dfrA1* via non-IS26 Tn7 transposon. Thus, strains like 206-05 and 210-04 (MIC >16/305 µg/mL) likely possess MGE-driven mechanisms robust enough to contribute to resistance, albeit less efficiently than truly resistant strains, while the sensitivity of strains like HI4320 have mechanisms still overcome by trimethoprim. Such prediction errors can be dangerous for treatment outcomes.

The genotype-phenotype discordance also highlights a significant limitation of current genomic prediction models: their reliance on curated databases of known resistance genes and mutations. Our study confirms that WGS-based prediction is highly reliable for well-characterized resistance determinants in *P. mirabilis* but can fail to predict phenotypes that are likely governed by cryptic mutations, efflux, and MGEs that impact gene expression, copy number, and functional redundancy. Not much has been reported on the mechanism of resistance involving efflux systems and membrane permeability [10, 83, 91]. Efflux systems such as *AcrAB-TolC* reported in many *P. mirabilis* strains can be overwhelmed by high drug concentrations, rather than the efficient enzymatic hydrolysis provided by a beta-lactamase and is linked to a very high multidrug-resistance rates [92]. Genotype based predictions are also prone to sequencing, assembly, and annotation errors, ideally needing long-read or reference-based sequencing to create hybrid assemblies. However, this can become impractical and costly for diagnostic laboratories.

Several limitations should be considered when interpreting our findings. First, the publicly available genomes were only derived from human hosts in the US, potentially biasing global population inferences. Second, our MGE analysis was restricted to the 22 clinical hybrid-assembled genomes to ensure accuracy, limiting the sample size for these comparisons but short reads are not an accurate predictor for deep MGE analysis. Third, while we identified compelling associations between MGE architecture in ST135 and AMR burden, it was done only in the sample set of 26 clinical isolate genomes and HI4320 with short-read assembly. Fourth, functional validation of IS26-mediated gene stacking and prophage transduction is necessary to confirm the mechanistic roles proposed here. Finally, our genotype-phenotype discordance analysis, though comprehensive, was necessarily limited by the number of clinical isolates with paired AST data.

In conclusion, while WGS has the potential to be used widely in clinical context for AMR predictions, our findings reinforce that assessing gene presence/absence does not replace phenotypic AST of *P. mirabilis* isolates for accurate clinical decision-making and surveillance. Accurate resistance prediction requires larger sampling size, database curation, resistance gene detection, and comprehensive characterization of the MGE landscape in *P. mirabilis*. These findings highlight the need for genomic surveillance targeting both high-risk lineages and the mobile elements that enable resistance gene flux across phylogenetic boundaries.

## Supporting information

Coding commands

## Acknowledgements

This work was supported by an American Heart Association (AHA) pre-doctoral fellowship 24PRE1195538 to ND, and by the National Institute of Diabetes and Digestive and Kidney Diseases under awards R01 DK123158 and R01 DK140371 to CEA.

## Author Contributions

ND and CEA designed experiments. ND, KC, and ALB conducted MIC experiments, ND and CEA analyzed results. ND performed genome sequencing, pangenome analysis, and AMR detection. BH and JW provided guidance on pangenome analysis, AMR detection, and data visualization. ND visualized the data and prepared the first draft of the manuscript. CEA provided project administration. All authors critically reviewed and revised the manuscript.

