## Supplemental Figures for "Pangenome Analysis of *Proteus mirabilis* Reveals Lineage-Specific Antimicrobial Resistance Profiles and Discordant Genotype-Phenotype Correlations"

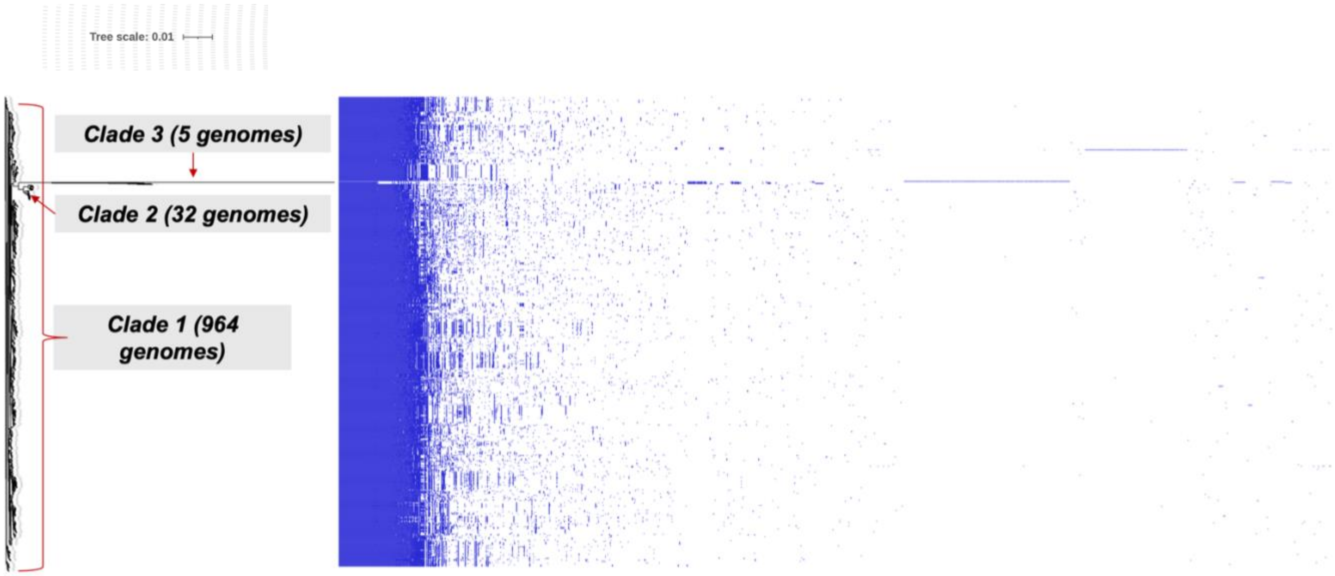


**Fig S1**. Mosaic pan genome with presence/absence matrix of core-genome phylogenetic tree for 1,001 *P. mirabilis* genomes. Three clades with different branch lengths (*sb1, sb2,* and *sb3*) are represented in the phylogenetic tree, where *sb2* has a clear difference in core genome.

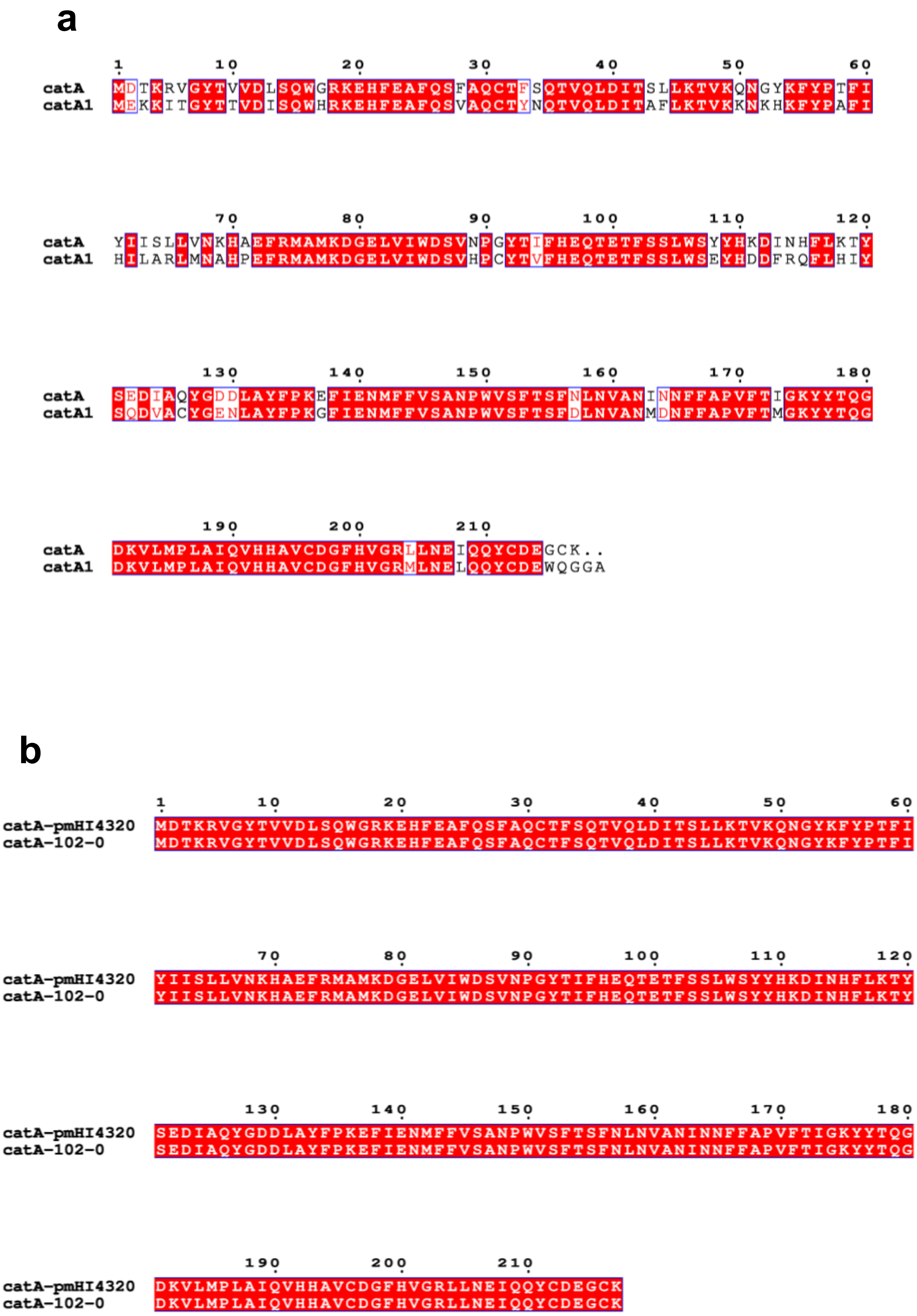
a
1 10 20 30 40 50 60
catA MDTKRVGYTVVDLSQWGRKEHFEAFQSFAQCTFSQTVQLDITSLLKTVKQNGYKFYPTFI
catA1 MEKKITGYTTVDISQWHRKEHFEAFQSVAQCTYNQTVQLDITAFLKTVKKNKHKFYPAFI
70 80 90 100 110 120
catA YIISLLVNKHAEFRMAMKDGELVIWDSVNPGYTIFHEQTETFSSLWSYYHKDINHFLKTY
catA1 HILARLMNAHPEFRMAMKDGELVIWDSVHPCYTVFHEQTETFSSLWSEYHDDFRQFLHIY
130 140 150 160 170 180
catA SEDIAQYGDDLAYFPKEFIENMFFVSANPWVSFTSFNLNVANINNFFAPVFTIGKYYTQG
catA1 SQDVACYGENLAYFPKGFIENMFFVSANPWVSFTSFDLNVANMDNFFAPVFTMGKYYTQG
190 200 210
catA DKVLMPLAIQVHHAVCDGFHVGRLLNEIQQYCDEGCK..
catA1 DKVLMPLAIQVHHAVCDGFHVGRMLNELQQYCDEWQGGA
b
1 10 20 30 40 50 60
catA-pmHI4320 MDTKRVGYTVVDLSQWGRKEHFEAFQSFAQCTFSQTVQLDITSLLKTVKQNGYKFYPTFI
catA-102-0 MDTKRVGYTVVDLSQWGRKEHFEAFQSFAQCTFSQTVQLDITSLLKTVKQNGYKFYPTFI
70 80 90 100 110 120
catA-pmHI4320 YIISLLVNKHAEFRMAMKDGELVIWDSVNPGYTIFHEQTETFSSLWSYYHKDINHFLKTY
catA-102-0 YIISLLVNKHAEFRMAMKDGELVIWDSVNPGYTIFHEQTETFSSLWSYYHKDINHFLKTY
130 140 150 160 170 180
catA-pmHI4320 SEDIAQYGDDLAYFPKEFIENMFFVSANPWVSFTSFNLNVANINNFFAPVFTIGKYYTQG
catA-102-0 SEDIAQYGDDLAYFPKEFIENMFFVSANPWVSFTSFNLNVANINNFFAPVFTIGKYYTQG
190 200 210
catA-pmHI4320 DKVLMPLAIQVHHAVCDGFHVGRLLNEIQQYCDEGCK
catA-102-0 DKVLMPLAIQVHHAVCDGFHVGRLLNEIQQYCDEGCK

**c**

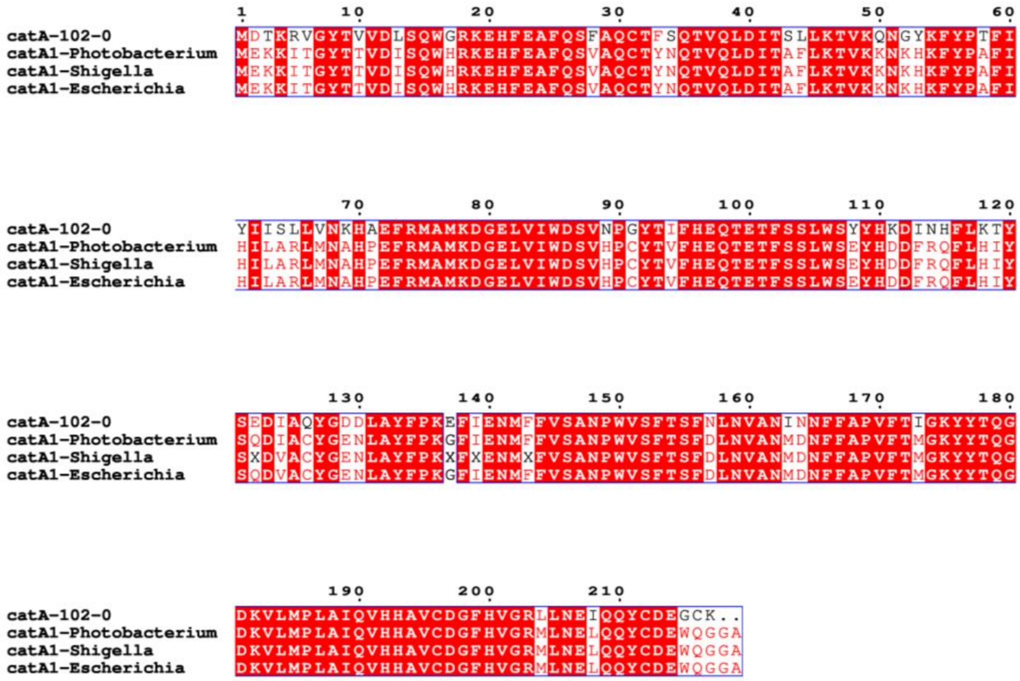


**Fig S2. Comparative sequence analysis of *catA* and *catA1* genes in *P. mirabilis* clinical isolates. a.** Amino acid equence comparison between *catA* and *catA1* demonstrates 78% amino acid sequence identity. **b.** Sequence comparison between the *catA* gene from strain 102-00 to the *catA* gene of reference strain *P. mirabilis* HI4320 shows 100% amino acid sequence identity. **c**. Sequence comparison between the *catA1* gene in different species of *Enterobacterales*.
