## Supplementary material for "Pangenome Analysis of *Proteus mirabilis* Reveals Lineage-Specific Antimicrobial Resistance Profiles and Discordant Genotype-Phenotype Correlations": Coding commands

### Supplementary Appendix: Tools and Databases

1. **FastQC v0.11.9-Java-11:** A high throughput sequence QC analysis tool.

```
fastqc "$INPUT_DIR"/*.fastq -o "$OUTPUT_DIR" -f fastq
```

2. **trimmomatic v0.39-Java-11.0.16:** Trimming ool for adapters from Illumina Sequencing.

```
java -jar $EBROOTTRIMMOMATIC/trimmomatic-0.39.jar PE -phred33 -threads 8
${infile} ${reverse_file} \
    ${base}_1.trim.fastq ${base}_1un.trim.fastq \
    ${base}_2.trim.fastq ${base}_2un.trim.fastq \
    ILLUMINACLIP:combined_adapters.fasta:2:30:10 LEADING:20 TRAILING:20
SLIDINGWINDOW:4:20 MINLEN:70
```

3. **spades v3.15.5:** Genome assembler.

```
spades.py --only-assembler --cov-cutoff auto --pe1-1 <r1file> --pe1-2 <r2file> -o
<output_dir>
```

4. **QUAST v5.2.0:** Quality Assessment Tool for Genome Assemblies.

```
quast.py <contigs_file> -o <sample_output> -m 500
```

5. **Unicycler v0.5.0:** An assembly pipeline for bacterial genomes-hybrid assembly.

```
unicycler -1 <r1_short_read_file> -2 <r2_short_read_file> -l <longread_file> -o
<output_dir> --spades_options "—memory ${memory mb}"
```

6. **flye v2.9.3:** Assembly- long + short read (hybrid).

```
flye --nano-raw "$ONT" --out-dir "$OUT_FLYE" --genome-size "$GENOME_SIZE" \
    &> "logs/${SAMPLE}. flye.log" || true
```

7. **bwa v0.7.17:** Index assembly and alignment tool.

```
bwa mem -t "$THREADS" "$ASM" "$R1" "$R2" >"logs/${SAMPLE}.bwa_mem.log"\

    | samtools view -@ "$THREADS" -bu - \
```

```
| samtools sort -@ "$THREADS" -o "$BAM" - || true
```

**samtools sort and BAM creation + index**

```
samtools sort -@ "$THREADS" -o "$BAM" -
  samtools index -@ "$THREADS" "$BAM" || true
```

8. **Pilon polishing; pilon v1.23-Java-11.0.16**

```
java -Xmx${JAVA_HEAP} -jar "$EBROOTPILON/pilon.jar" --genome "$ASM" --bam "$BAM"
--changes --vcf --output "polished_${SAMPLE}" --outdir "$OUT_PILON" \
    &> "${OUT_PILON}/pilon.log" || true
```

9. **Prokka v1.14.5**

```
prokka --compliant --cpus 16 --prefix ${prefix} --outdir ${output_dir} --force
${infile}
```

10. **PubMLST database:** Open-access, curated databases that integrates population sequence data with provenance and phenotypes. https://pubmlst.org/

```
mlst --scheme proteus_spp "$FASTA_FILE" > "$MLST_DIR/${SAMPLE_NAME}_mlst.csv"
```

11. **roary v3.13.0**

```
roary -i 90 -e --mafft -f *.gff
```

12. **FastTree v2.1.11**

```
FastTree -gtr -gamma -nt core_gene_alignment.aln > SampleID_tree.nwk
```

13. **Raxml-ng/1.2.1**

```
raxml-ng --all --msa core_gene_alignment.aln --model GTR+G --prefix ${prefix}--
bs-metric fbp,tbe
```

14. **AMRFinderPlus v 3.11.18**

```
amrfinder --nucleotide <NUC_FASTA> -u
```

15. **ABRicate v1.0.1:**

```
abricate -db <db_name> file.fasta > results.tsv
```

16. **ISEScan v1.7.3**

Dependencies:
FragGeneScan1.3
hmmer
LIBRARY:ISEScan-1.7.3/ssw201507

```
python3 ISEScan-1.7.3/isescan.py --seqfile "$fasta" –output
"IS_scan_results_NCBI/${sample}" --nthread "$THREADS"
```

17. **PHASTER** (PHAge Search Tool Enhanced Release): web server for the rapid identification and annotation of prophage sequences within bacterial genomes and plasmids.

https://phaster.ca/

18. **ICEberg 3.0:** Web server for predicting the conjugative and accessory modules.

https://tool2-mml.sjtu.edu.cn/ICEberg3/ICEfinder.php

19. **R v4.3.0**
20. **Stata/IC version 15.1**
